# The assembly landscape of the complete B-repeat superdomain from *Staphylococcus epidermidis* strain 1457

**DOI:** 10.1101/2023.01.04.522776

**Authors:** Alexander E. Yarawsky, Andrew B. Herr

**Affiliations:** Division of Immunobiology, Cincinnati Children’s Hospital Medical Center, Cincinnati, Ohio, USA; Division of Infectious Diseases, Cincinnati Children’s Hospital Medical Center, Cincinnati, Ohio, USA; Department of Pediatrics, University of Cincinnati College of Medicine, Cincinnati, Ohio, USA

**Keywords:** analytical ultracentrifugation, protein self-assembly, equilibrium, biofilm formation, amyloid, circular dichroism, dynamic light scattering, metal-binding protein

## Abstract

The accumulation-associated protein (Aap) is the primary determinant of *Staphylococcus epidermidis* device-related infections. The B-repeat superdomain is responsible for intercellular adhesion that leads to the development of biofilms occurring in such infections. It was recently demonstrated that Zn-induced B-repeat assembly leads to formation of functional amyloid fibrils, which offer strength and stability to the biofilm. Rigorous biophysical studies of Aap B-repeats from *S. epidermidis* strain RP62A revealed Zn-induced assembly into a stable, reversible dimer and tetramer, prior to aggregation into amyloid fibrils. Genetic manipulation is not tractable for many *S. epidermidis* strains, including RP62A; instead, many genetic studies have used strain 1457. Therefore, to better connect findings from biophysical and structural studies of B-repeats to *in vivo* studies, the B-repeat superdomain from strain 1457 was examined. Differences between the B-repeats from strain RP62A and 1457 include the number of B-repeats, which has been shown to play a critical role in assembly into amyloid fibrils, as well as the distribution of consensus and variant B-repeat subtypes, which differ in assembly competency and thermal stability. Detailed investigation of the Zn-induced assembly of the full B-repeat supderdomain from strain 1457 was conducted using analytical ultracentrifugation. Whereas the previous construct from RP62A (Brpt5.5) formed a stable tetramer prior to aggregation, Brpt6.5 from 1457 forms extremely large stable species on the order of 28- and 32-mers, prior to aggregation into similar amyloid fibrils. Importantly, both assembly pathways proceed through the same mechanism of dimerization and tetramerization, and both conclude with the formation of amyloid-like fibrils. Theoretical discussions of the energy landscape of B-repeats from different strains and of different length are provided with considerations of biological implications.

**Statement of Significance:** *Staphylococcus epidermidis* is a major pathogen responsible for device-related infections. The primary factor responsible for such infections is the accumulation-associated protein, Aap, through its ability to mediate formation of dense surface-adherent communities of bacteria known as biofilms. Our lab recently demonstrated that the B-repeat superdomain of Aap from strain RP62A undergoes Zn-dependent assembly to form functional amyloid fibrils that improve the strength and resilience of biofilms *in vitro*. These amyloid fibrils may be responsible for the difficulty in treating device-related infections. However, strain 1457 is commonly used for genetic manipulation. In this manuscript, the Zn-dependent assembly of B-repeats from strain 1457 is shown to lead to the same outcome of amyloidogenesis, although it occurs through different intermediate oligomeric states.

## INTRODUCTION

*Staphylococcus epidermidis* has long been recognized as one of the most problematic organisms in medical device-related infections (1–4). *S. epidermidis* is a human commensal that is abundant on the skin but can also contaminate devices such as catheters, joint replacements, and even pacemakers after surgical implantation. Following bacterial adherence to a surface, the bacteria adhere to one another to form a resilient cluster known as a biofilm (3). While cell-to-cell accumulation can be mediated via production of the polysaccharide known as poly-*N*-acetylglucosamine (PNAG), data from a study of clinical isolates show the presence of the accumulation-associated protein (Aap) in 89% of prosthetic hip and knee joint infections (5). Furthermore, a recent study using a rat catheter model demonstrated that Aap is required for infection (6).

The cell-wall anchored Aap, regardless of strain, has a well-conserved arrangement of domains (6,7). At the N-terminus, a series of short A-repeats with unknown function is present, followed by a lectin domain responsible for surface attachment (8–10) and SepA proteolytic cleavage sites on either side of the lectin domain (11). Downstream of the SepA site is the B-repeat superdomain containing 5-17 B-repeats – each composed of a G5 subdomain and spacer (also called E) subdomain. SepA cleavage occurs during biofilm formation, exposing the B-repeat superdomain and allowing for cell-to-cell adhesion via Zn^2+^-mediated B-repeat interactions (11–13). The B-repeat superdomain protrudes outward from the cell surface due to a highly extended stalk region that is rich in proline and glycine (14,15). At the C-terminus, a LPXTG Sortase A motif is present, which anchors Aap to the peptidoglycan layer of the bacterial cell wall (16,17).

Extensive biophysical and structural work has been performed on Aap B-repeats from *S. epidermidis* strain RP62A (7,13,18–22). However, much of the functional *in vivo* work has been performed using strain 1457, given it is more amenable to genetic manipulation (6,23). The goal of this manuscript is to compare B-repeat assembly and amyloidogenesis of Aap B-repeats from strain 1457 to previous analyses of B-repeats from strain RP62A, such that results from biophysical and structural studies can be translated more appropriately to B-repeats from other strains and to plan future genetic manipulations of *S. epidermidis* to rigorously test the impact of B-repeat assembly and amyloidogenesis.

Specifically, we begin by examining the assembly of wild-type Brpt6.5 from strain 1457 in the presence of Zn^2+^. Interestingly, an extremely large stable oligomer is observed. As with Brpt3.5 and Brpt5.5 from RP62A Aap, Brpt6.5 also assembles into functional amyloid-like fibrils (21). To examine dimerization, a Brpt6.5 mutant with the E203 position of each B-repeat mutated to alanine was used. Consistent with RP62A Aap Brpt1.5 crystallography and biophysical data (18), the Brpt6.5 7xE203A mutant was unable to dimerize in the presence of Zn – indicating similar mechanisms of dimerization. The RP62A Aap Brpt5.5 tetramer was recently identified as a critical step in assembly and aggregation into fibrils via the H85 position of each B-repeat (20). Similarly, H85A mutations in Brpt6.5 could prevent tetramer formation or allow tetramer formation but preclude amyloidogenesis. In conclusion, the B-repeats from Aap from strain 1457 utilize similar mechanisms of assembly and aggregation, with significant differences in early assembly equilibrium constants and the stability of intermediates on the way to the terminal amyloid-like fibrils.

## MATERIALS AND METHODS

### Protein Design

The Brpt6.5 WT, 6xH85A, and 7xE203A constructs were synthesized by Life Technologies, Inc., GeneArt® and subcloned into the pENTR vector. The 6xH85A mutant contained the following mutations: H85A, H213A, H341A, H469A, H597A, and H725A. The 7xE203A mutant contained the following mutations: H75A, H203A, H331A, H459A, H587A, H715A, E843A. The VC (**v**ariant sequence mutated to **c**onsensus sequence) construct was also synthesized by Life Technologies, Inc., GeneArt® and subcloned into the pENTR vector. The VC mutations included N789D, D791N, K793A, E796T, R798K, and K800V. Residues V842 and E843 are already present in the WT sequence, so no sequence modification of these two residues was necessary for the VC construct. Each construct contained a tobacco etch virus (TEV) protease cleavage site directly upstream of the gene. At the C-terminus, an added glycine was followed by another TEV protease cleavage site (ENLYFQ|G) and Strep-tag (WSHPQFEK) using the Agilent QuikChange II Site-Directed Mutagenesis Kit. Single H85A mutants of Brpt6.5 were produced from the parent construct using QuikChange mutagenesis as was previously done for Brpt5.5 (20), with the exception that here the alanine residues from the 6xH85A mutant were being mutated back to histidine. The LR Clonase reaction was used to transfer the gene of interest and added Strep-tag to a destination vector containing an N-terminal His6-MBP tag. Aap Brpt6.5 from strain 1457 contained amino acids 608-1454 from UniProt Entry: A0A075IHN3. Aap Brpt5.5 from strain RP62A contained amino acids 1505-2223 from UniProt Entry: Q5HKE8. B-repeat constructs used in this study are summarized in Figure S1.

### Protein Expression and Purification

Brpt6.5 constructs were expressed in BLR(DE3) cells grown to an OD_600_ of ~0.8 at 37 °C, shaking, in 1 L of LB media containing ampicillin and tetracycline antibiotics to maintain the cell line and plasmid. Then, the cultures were chilled to 10 °C in an ice bath. Ethanol was added to a final concentration of 2%, and protein production was induced with 200 μM IPTG. Cultures were incubated at 20 °C overnight, with shaking. The cultures were centrifuged to pellet the bacteria, which were then resuspended in 20 mM Tris pH 7.4, 300 mM NaCl with a protease inhibitor cocktail tablet (Roche). Resuspended pellets were stored at −80 °C.

Pellets were lysed by sonication. The lysate was then centrifuged, and the supernatant was filtered using 0.45 μm and 0.20 μm bottle-top filters. Protein was purified using a 20 mL NiNTA column (Cytiva) attached to an Akta M FPLC (GE Healthcare) and with a 300 mM imidazole elution buffer. The fractions containing eluted protein were pooled and dialyzed into 20 mM Tris pH 7.4, 150 mM NaCl overnight at 4 °C before being loaded onto a 5 mL StrepTrap HP column (Cytiva) and eluted with 2.5 mM desthiobiotin (Sigma). The pooled elution fractions were then incubated at room temperature, rocking or stirring gently, with TEV protease for approximately 18-24 hours – adding additional protease after the first 6-16 hours. The protein solution was then filtered and run over a 20 mL NiNTA column – collecting the flow-through, followed by the 5 mL StrepTrap HP column – again collecting the flow-through. When deemed necessary based on SDS-PAGE or AUC, an additional purification step using a 24 mL Superose 6 size exclusion column (Cytiva) was implemented.

### Circular Dichroism

CD experiments were performed on an Aviv 215 CD spectrophotometer with a Peltier junction temperature control system. Far-UV wavelength scans were performed using a 0.1 mm quartz cuvette at 20 °C, with 5 scans being averaged, and a 3 second averaging time at each wavelength step. Protein was at 1.5 mg/ml in 50 mM MOPS pH 7.2, 50 mM NaCl. The conversion from machine units (millidegrees, θ) to mean residues ellipticity, [θ] – degrees cm^2^ dmol^-1^ residue^-1^, was performed using Equation (1):

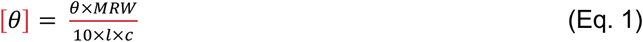

MRW is the mean residue weight for WT (107.4 g mol^-1^ residue^-1^), VC (107.2 g mol^-1^ residue^-1^), 7xE203A (106.9 g mol^-1^ residue^-1^), or 6xH85A (106.9 g mol^-1^ residue^-1^). The pathlength, *l*, was in cm units and concentration in mg/ml units.

Thermal denaturation experiments were performed in the same cuvette and buffer, and at the same protein concentration. The experiments followed the signal at 205 nm from 20 °C to 80 °C, in 1 °C steps. The averaging time at each step was 10 seconds. Where direct comparisons are made in the figures, each sample was dialyzed into the same buffer to ensure accurate comparisons were made, as pH and salt concentration differences may cause significant effects on thermal stability. Thermal denaturation experiments were analyzed in SigmaPlot 12.5 (Systat Software, Inc.) using a two-state model correcting for pre- and post-transition linear changes (24). It was assumed that there was no change in heat capacity. Data are presented as fraction folded, according to Equation (2):

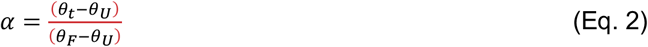

where θ_t_ is the MRE at a given temperature, θ_F_ is the MRE of the folded monomer, θ_U_ is the MRE of the unfolded fold (24).

Temperature-dependent wavelength experiments were carried out with 1.8 mg/ml protein in 50 mM MOPS pH 7.2, 50 mM NaCl with 5 mM ZnCl_2_ added directly to the sample. Samples were examined in a 0.5 mm cuvette. A macro was used to run experiments sequentially without any time manual input. Scans were single replicates with 3 second averaging time at each wavelength from 300 nm to 190 nm, at 10 °C increments from 20 °C to 80 °C, then another scan after cooling to 20 °C.

Temperature-dependent aggregation experiments were performed as previously described (20), but with 0.50 mg/ml Brpt6.5. Prior to the start of the experiment, 3.50 mM ZnCl_2_ was added directly to the sample. A wavelength of 225 nm was used to monitor formation of amyloid-like aggregate formed by WT, and a wavelength of 218 nm was used to follow unfolding of the Brpt6.5 6xH85A mutant.

### Sedimentation Velocity Analytical Ultracentrifugation

AUC experiments were performed using a Beckman Coulter, Inc. ProteomeLab XL-I. Unless otherwise specified, experiments were run at 48,000 rpm using Spin Analytical 1.2 cm meniscus-matching centerpieces at 20 °C. To match menisci, a short (~5-10 min) spin at 3,000 – 5,000 rpm was used while monitoring the fringe display window to ensure successful transfer of buffer to the protein sector. The cells were removed from the rotor, gently rocked by hand to redistribute the protein gradient, then carefully re-aligned in the rotor. The AUC was allowed to equilibrate at least until the temperature display read 20.0 °C for several minutes. The experiments were allowed to run overnight – usually at least 16 hours, or until sedimentation of protein appeared complete. Sedimentation velocity data were analyzed by WDA (wide distribution analysis) within SEDANAL (25,26) or using the c(s) distribution model within SEDFIT (27).

The relationship between the sedimentation coefficient, frictional ratio, and molecular weight (M) was used to estimate the sedimentation coefficient of Brpt6.5 oligomers (Equation (3)):

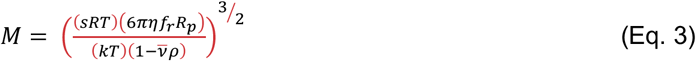

where *s* is the sedimentation coefficient (s x 10^-13^ sec), *f_r_* is the frictional ratio, *R_p_* is the radius of an equivalent sphere calculated according to Teller (28) (6.72 x 10^-9^ * M_r_^(1/3)^). *R* is the molar gas constant (J/(mol K)), *k* is the Boltzmann constant (1.381 x 10^23^ J/K). *T* is the temperature in Kelvin, *η* is the viscosity (poise), *ρ* is the density (g/ml), and 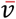 is the partial specific volume of the protein, estimated from the amino acid sequence using SEDNTERP v3.0.3 (29).

SEDNTERP v3.0.3 was also used to determine the hydrodynamic nonideality coefficient (*k_s_*) according to Equation (4):

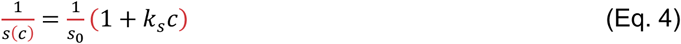

Where s(c) is the concentration dependence of the sedimentation coefficient, s_0_ is the sedimentation coefficient at zero concentration, c is the concentration, and *k_s_* is the hydrodynamic nonideality coefficient. The values for *s_0_*, *k_s_*, and error estimates are listed in Table S1.

### Sedimentation Equilibrium Analytical Ultracentrifugation

Sedimentation equilibrium data were collected at 4,000 rpm using absorbance optics at 233 nm, 240 nm, and 277 nm. Six-channel centerpieces were used. Loading concentrations were approximately 0.5, 0.3, and 0.1 mg/ml Brpt6.5 WT in 50 mM MOPS pH7.2, 50 mM NaCl, 3.50 mM ZnCl_2_. Equilibrium was verified by the MATCH utility within HeteroAnalysis (30). Data were analyzed within SEDANAL (31) using a model containing a single, ideal species or with the addition of a monomer with a fixed MW.

### Dynamic Light Scattering

DLS experiments used a Malvern Zen 3600 Zetasizer Nano and a low-volume quartz cuvette. Brpt6.5 samples were adjusted to 0.50 mg/ml in 50 mM MOPS pH 7.2, 50 mM NaCl, filtered using a 0.2 μm syringe filter, and had 3.50 mM of filtered ZnCl_2_ added just prior to the experiment. The DLS method took 2 °C temperature steps with 120 seconds of equilibration time. Each measurement was made in triplicate with automatic measurement duration. Changes in viscosity were considered at each temperature step in the Zetasizer program.

### Turbidity Assay

Brpt6.5 samples were dialyzed into the same buffer preparation of 50 mM MOPS pH 7.2, 50 mM NaCl and adjusted to 0.50 mg/ml. Triplicate samples of protein and dialysis buffer were prepared. A BioMate 3S UV-Vis spectrophotometer was set to the multiwavelength mode (280 nm, 400 nm, and 700 nm) and blanked with dialysis buffer. A 200 μl sample was transferred to a quartz low-volume cuvette and a scan was taken to ensure expected baseline readings near 0.000 at 400 nm and 700 nm, with a 280 nm reading indicative of 0.5 mg/ml protein (or zero for buffer). Then, thirty 1-μl aliquots of a 500 mM ZnCl_2_ solution were added to the sample, with gentle shaking between each addition and scan. The spectrophotometer measurements were taken at room temperature. This resulted in a final ZnCl_2_ concentration of approximately 65 mM. The data collected at 400 nm are presented, as these values were within the linear range of the spectrophotometer (<1.0 OD) and showed greater signal than that observed at 700 nm, without any ZnCl_2_ absorbance that was observed at 280 nm.

### Thioflavin T Assay

Samples of Brpt6.5 WT dialyzed into 50 mM MOPS pH 7.2, 50 mM NaCl were added to wells of a 96-well plate at a concentration of 4 mg/ml. Four replicate wells were set up for each condition. Data presented were collected after 24 hours of shaking at 37 °C at 200 rpm, with the plate sealed. Thioflavin T (ThT) was added to a final concentration of 20 μM at the start of the experiment. The total volume of each well was 100 μl. The plates were scanned using a Biotek Synergy plate reader with an excitation wavelength of 440 nm and emission wavelength of 482 nm, using the bottom optics position and a gain of 100. To test for significance, a Student’s *t* test was performed in MS Excel (two-tailed, two-sample unequal variance). Significance was identified when *p* < 0.05.

### Transmission Electron Microscopy

A 5-μl sample was taken from the ThT assay at the 24-hour timepoint. The sample was added to a 200 mesh Formvar carbon/copper grid. After 2 min, the grid was washed with 400 μl of Milli-Q water, and Whatman filter paper was used to wick the liquid off the grid. The grid was then stained for 30 seconds using 1% uranyl acetate that was added dropwise. The grid was then washed again with 400 μl of Milli-Q Water before being dried with filter paper and left to air-dry at least overnight. Images were captured using an AMT 2k CCD camera on a Hitachi 7600 microscope at an accelerating voltage of 80 kV.

## RESULTS

### A comparison of Aap from strain 1457 and RP62A

The overall domain arrangement of Aap is conserved across strain 1457 and RP62A - as shown in Figure 1A and Figure S2. The primary differences between these and other strains are the number of B-repeats (6,7). In addition, the sequence of each B-repeat may be classified as a “consensus” or “variant” subtype. In the variant repeats, there are eight residues in the G5 subdomain of the B-repeat that replace those found in the consensus (Figure 1B (purple), Figure S1). Based on previous studies using Brpt1.5 constructs, it was found that consensus B-repeats were better able to assemble in the presence of Zn^2+^, while variant B-repeats conferred higher thermal stability of the folded monomer (7). Brpt6.5 from strain 1457 Aap is composed of 6 full consensus repeats, with the C-terminal half-repeat (G5 subdomain) being the variant subtype. Interestingly, Shelton, et al. observed a significant position-dependent effect of the variant sequence in the context of Brpt1.5, with the C-terminal G5 subdomain being the major determinant of Zn^2+^-dependent assembly and thermal stability (7). Thus, while Brpt6.5 (1457) and Brpt5.5 (RP62A) are both predominantly consensus, the presence of a variant B-repeat in the C-terminal half-repeat position in Aap from 1457 could potentially have major consequences on assembly and stability.

**Figure 1.**
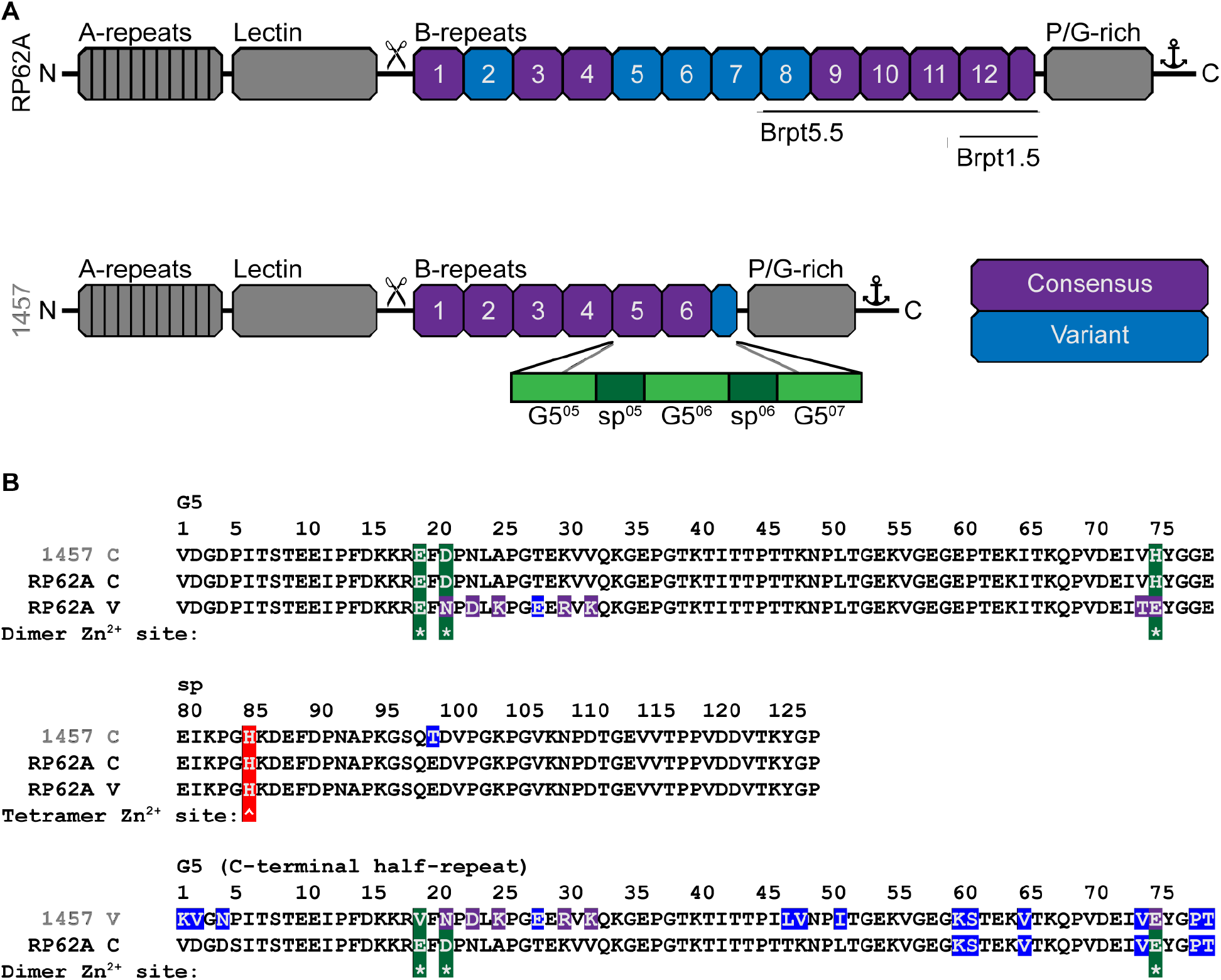
A comparison of Aap from *S. epidermidis* strain RP62A and 1457. (A) Domain arrangements of Aap from each strain. B-repeats are colored according to whether the sequence cassette is consensus or variant, as described in the text (7). (B) Examples of the consensus and variant B-repeats are shown. Residues highlighted in purple represent the residues present in the variant cassette. Aap from strain 1457 only contains a variant cassette in the C-terminal half-repeat. The C-terminal half-repeat from Aap from strain RP62A is consensus. In both cases, the C-terminal half-repeat has lesser identity to the other G5 subdomains of Aap (highlighted in blue). Residue positions found to coordinate Zn^2+^ during dimerization are marked with an asterisk (*) and highlighted in green, while residue H85 is marked with a caret (^) and highlighted in red to denote its involvement in tetramerization.

### Brpt6.5 secondary structure and stability

B-repeats from strain RP62A Aap exhibit β-sheet and random coil content, regardless of the G5 sequence type (consensus or variant) or the number of B-repeats in the construct (7,13,18,20,21). The Far-UV CD spectrum of Brpt6.5 WT monomer (without Zn^2+^ present) was comparable to those observed previously for RP62A Aap B-repeats (Figure 2A). Thermal denaturation data were fit to a two-state transition (monomer and unfolded monomer). Brpt6.5 WT showed a Tm of 55.1 °C (Figure 2B, Table 1), with a clear loss of β-sheet signal evident at 80 °C relating to the fully unfolded state (Figure 2A). Additional constructs (Brpt6.5 7xE203A and Brpt6.5 6xH85A) were produced to analyze the mechanisms of Brpt6.5 oligomerization (described below). The Brpt6.5 7xE203A mutant contained similar secondary structure content and similar folding stability compared to WT (Figure 2, Table 1). The Brpt6.5 6xH85A also showed similar secondary structure content (Figure 2A) but showed a significant decrease in Tm from 55.1 °C to 50.5 °C (Figure 2B). This is consistent with observations for Brpt5.5 5xH85A mutant from strain RP62A, as the H85 residue in each B-repeat is involved in a hydrogen-bonding network (20,22). Overall, the WT, 7xE203A, and 6xH85A constructs are properly folded, consistent with the expected secondary structure content, and are thermodynamically stable up to 50-55 °C.

**Figure 2.**
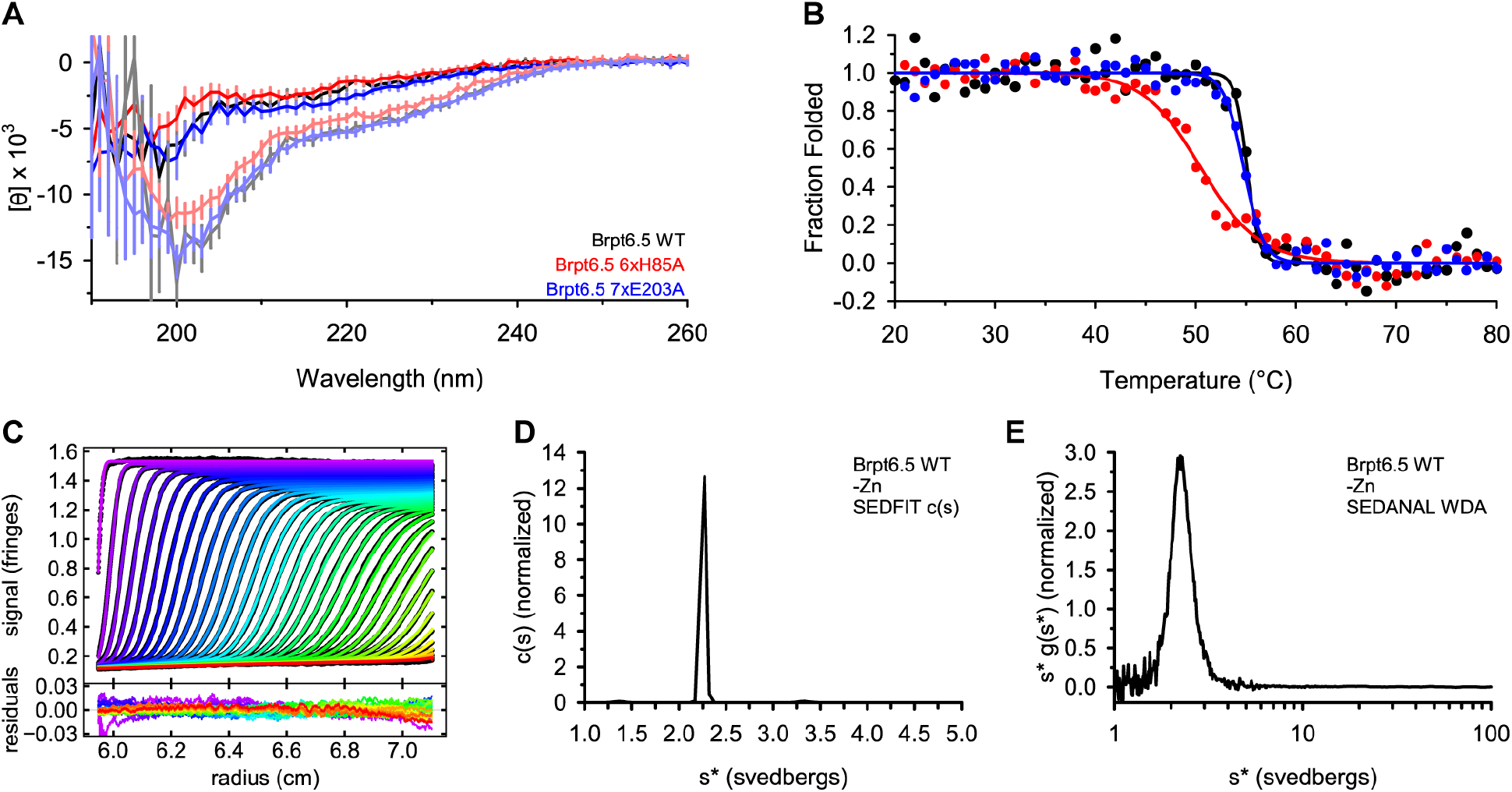
Brpt6.5 folding and behavior in solution without Zn^2+^. (A) Far-UV CD wavelength scans showing similar secondary structure content across all constructs. Dark colored lines were obtained at 20 °C, while light colored lines were obtained at 80 °C. (B) Thermal denaturation CD experiments monitoring the CD signal at 205 nm. Each dataset was fit to a two-state model. The markers indicate data points, while the solid lines show the best fit. Brpt6.5 6xH85A shows a significant decrease in thermal stability. Data are shown as fraction folded. CD data were collected in 50 mM MOPS pH 7.2, 50 mM NaCl, at 1.5 mg/ml protein. (C) Sedimentation velocity boundaries of Brpt6.5 WT at 48,000 rpm without ZnCl_2_. The SEDFIT c(s) fit lines are colored from purple (early time scans) to red (late time scans). Data are shown as black markers behind the fit lines, with residuals shown in the bottom panel. The sedimentation velocity data were analyzed by SEDFIT c(s) (D) and SEDANAL WDA (E). Data were collected in 50 mM MOPS pH 7.2, 50 mM NaCl, at 0.5 mg/ml protein.

**Table 1.** CD thermal denaturation results.

| Sample | $T_m$ (°C) | $\Delta H$ (kcal/mol) |
| --- | --- | --- |
| WT | $55.1 \pm 0.2$ | $-360 \pm 85$ |
| 7xE203A | $54.8 \pm 0.2$ | $-220 \pm 29$ |
| 6xH85A | $50.5 \pm 0.5$ | $-72 \pm 10$ |

\*Determined at 1.5 mg/ml Brpt6.5 WT in 50 mM MOPS pH 7.2, 50 mM NaCl in a 0.1 mm cuvette.

### Zn^2+^-induced assembly of Brpt6.5 WT

In the context of Aap from RP62A, a major function of the B-repeat superdomain is to mediate intercellular interaction of adjacent cells in the developing biofilm. This occurs via a Zn^2+^-dependent mechanism that includes dimer and tetramer assembly states. The tetramer state can then undergo a conformational change that triggers aggregation into functional amyloid fibrils, which apparently function to further stabilize the biofilm. The *S. aureus* ortholog of Aap, known as SasG, has also been shown to use Zn^2+^ in its role in intercellular adhesion (32). The relevance of these Zn^2+^-dependent interactions was confirmed in biofilms via the ability for DTPA (a metal chelator) to inhibit or disrupt biofilm formation in *S. epidermidis* RP62A expressing Aap or *S. aureus* USA300 expressing SasG (13,21). Zn^2+^-dependent heterophilic interaction between Aap from strain 1457 and SasG from *S. aureus* strain SH1000 has been shown via single-cell force measurements (32). Based on this observation, Aap from strain 1457 is likely to mediate intercellular adhesion in biofilms via the same Zn^2+^-dependent mechanism as described for strain RP62A (13,21).

The ability of the WT Brpt6.5 construct from strain 1457 Aap to assemble in the presence of Zn^2+^ was evaluated using sedimentation velocity analytical ultracentrifugation (AUC). In the absence of Zn^2+^, Brpt6.5 WT sedimented relatively slowly at 48,000 rpm (Figure 2), with a sedimentation coefficient (2.26 S) only slightly higher than that of Brpt5.5 (2.20 S (20)) and an estimated molecular weight near that of the sequence-based MW. Brpt6.5, like Brpt5.5, has an unusually large frictional ratio due to the extended conformation of the B-repeats (*f/f*_0_ = 3.48; Table 2). In the presence of Zn^2+^, Brpt6.5 WT showed one or more rapidly-sedimenting species which had not been observed previously for other B-repeat constructs from strain RP62A Aap (Figure 3). To determine the best experimental conditions with which to collect data on this system, a series of velocity experiments were run on Brpt6.5 + 3.50 mM ZnCl_2_ at speeds ranging from 18,000 rpm to 60,000 rpm. At the two lower speeds, many scans can be collected for the faster-sedimenting species, however, the slower-sedimenting species (monomer) does not completely sediment (Figure 3A and Figure 3B). At higher speeds the monomer fully sediments, but there is less time for data collection before the larger species sediment (Figure 3C and Figure 3D). Analysis via SEDFIT c(s) (Figure 3E) is robust in detection of the monomer (~2 S), while SEDANAL WDA (Figure 3F) is less able to provide consistent details on the monomer due to the incomplete sedimentation at lower speeds. Both approaches show strong signal relating to the ~26 S species, with the SEDFIT c(s) distribution showing two reaction boundaries ~26 S at higher speeds. This observation indicates that there may be more than one oligomer sedimenting near ~26 S.

**Figure 3.**
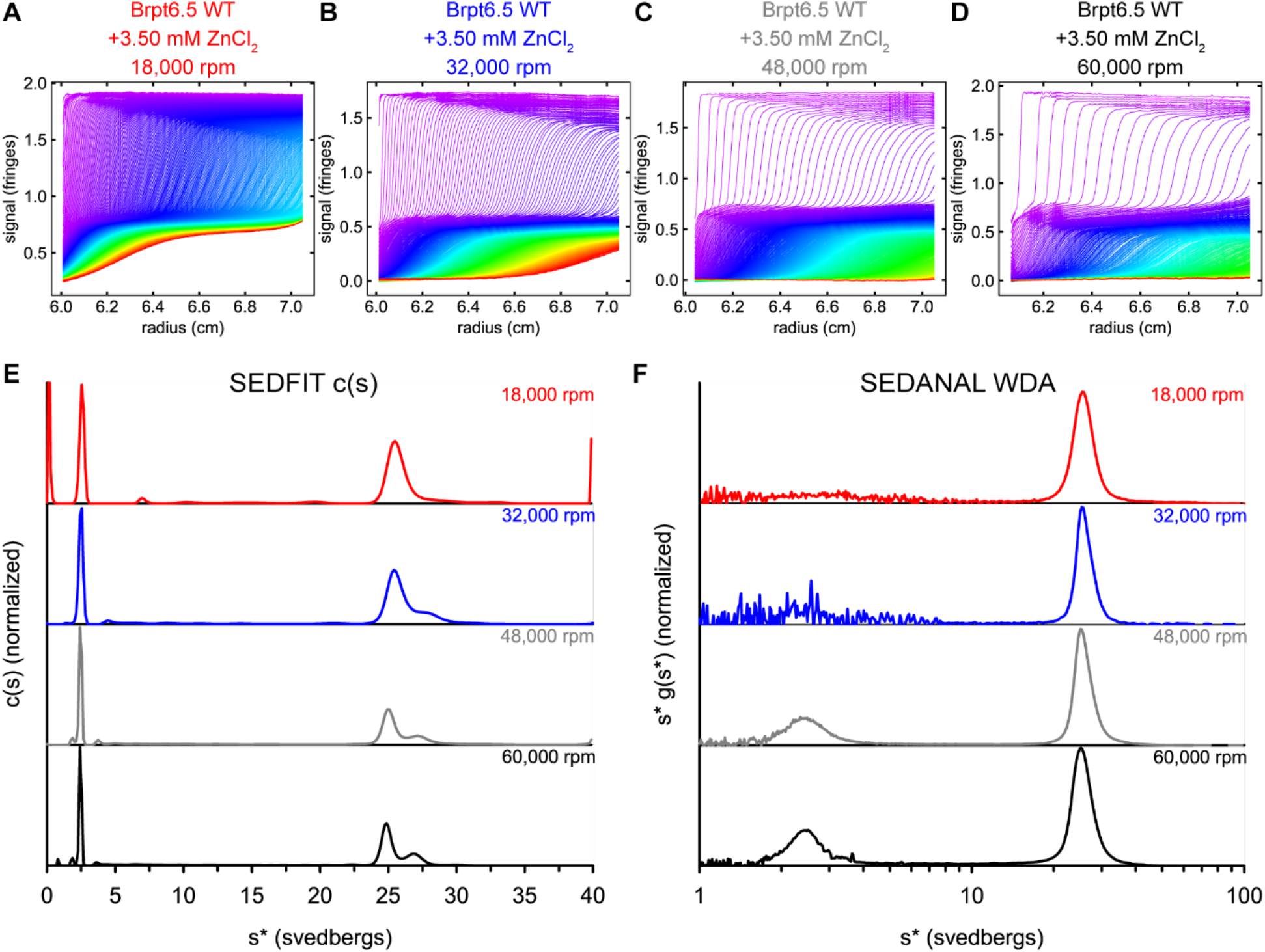
Brpt6.5 forms large oligomers in the presence of Zn^2+^. Sedimentation boundaries collected during sedimentation velocity AUC experiments of Brpt6.5 WT in the presence of 3.50 mM ZnCl_2_ at (A) 18,000 rpm, (B) 32,000 rpm, (C) 48,000 rpm, and (D) 60,000 rpm. Data presented were collected using interference optics. The colors indicate time of data collection, from early time points (purple) to late time points (red). Panels (A) - (D) were generated using GUSSI (36) after SEDFIT c(s) analysis. Both the TI and RI noise was removed. The datasets shown in (A) – (D) were analyzed by (E) SEDFIT c(s) analysis (27) or (F) SEDANAL WDA (25,26).

**Table 2.** Sedimentation velocity AUC results.

| Analysis | $S_w$ | $f/f_0$ | MW |
| --- | --- | --- | --- |
| SEDFIT c(s) | 2.26 | 3.48 | 103 kDa |
| SEDANAL WDA | $2.33 \pm 0.04$ | | |

\*Determined at 0.5 mg/ml Brpt6.5 WT in 50 mM MOPS pH 7.2, 50 mM NaCl. Expected MW based on sequence is 91.7 kDa.

A series of experiments at increasing Zn^2+^ concentrations was conducted to test the hypothesis that there may be multiple oligomers sedimenting near ~26 S. Firstly, Brpt6.5 WT was evaluated in the absence of Zn^2+^ across a range of protein concentrations. The lack of shift in the c(s) distribution toward higher s* (Figure 4A) and the decrease in s* with increasing concentration (Figure 4C) indicates that Brpt6.5 WT does not assemble in the absence of Zn^2+^ but shows hydrodynamic nonideality. This is consistent with previous reports for RP62A Aap Brpt1.5 and Brpt5.5 (7,13,20). A series of panels shows the c(s) distributions observed when Brpt6.5 WT was dialyzed into buffer containing increasing amounts of ZnCl_2_ (Figure 4D, 4E, black lines). A reaction boundary was observed near 2 S in the absence of Zn^2+^, consistent with the monomeric species in the absorbance measurements discussed above (Figure 2). As the ZnCl_2_ concentration was increased to 1.75 mM, the emergence of a reaction boundary near 26 S was detectable. This reaction boundary was observed up to the highest ZnCl_2_ concentration tested, with notable asymmetry and a shift in the S value, both of which are suggestive that a reaction boundary, not a discrete species, is observed. This is to be expected due to the rapid kinetics of assembly, as previously observed for B-repeats (7,13,18–22,33). Interestingly, in the lower S-value range, the monomer species is predominant until 6.5 mM ZnCl_2_, at which point a broader distribution is observed up to ~7 S, where the Brpt5.5 tetramer sediments (20,22). A clear trend is observed involving the weight-averaged sedimentation coefficient (sw) over the entire c(s) distribution and the ZnCl_2_ concentration (Figure 4F). The sw will be affected by the protein concentration, which may be decreased due to aggregation at higher Zn^2+^ concentrations. Therefore, Figure 4G shows the concentration of protein observed in the AUC experiment and is based on the area under the c(s) distributions. No major loss of WT protein was observed until 8 mM ZnCl_2_, at which point aggregation was visible after dialysis. Overall, the ~26 S species seem to be stable intermediates on the pathway to amyloidogenesis, rather than a transient, heterogeneous set of aggregates or an isodesmic assembly (34,35).

**Figure 4.**
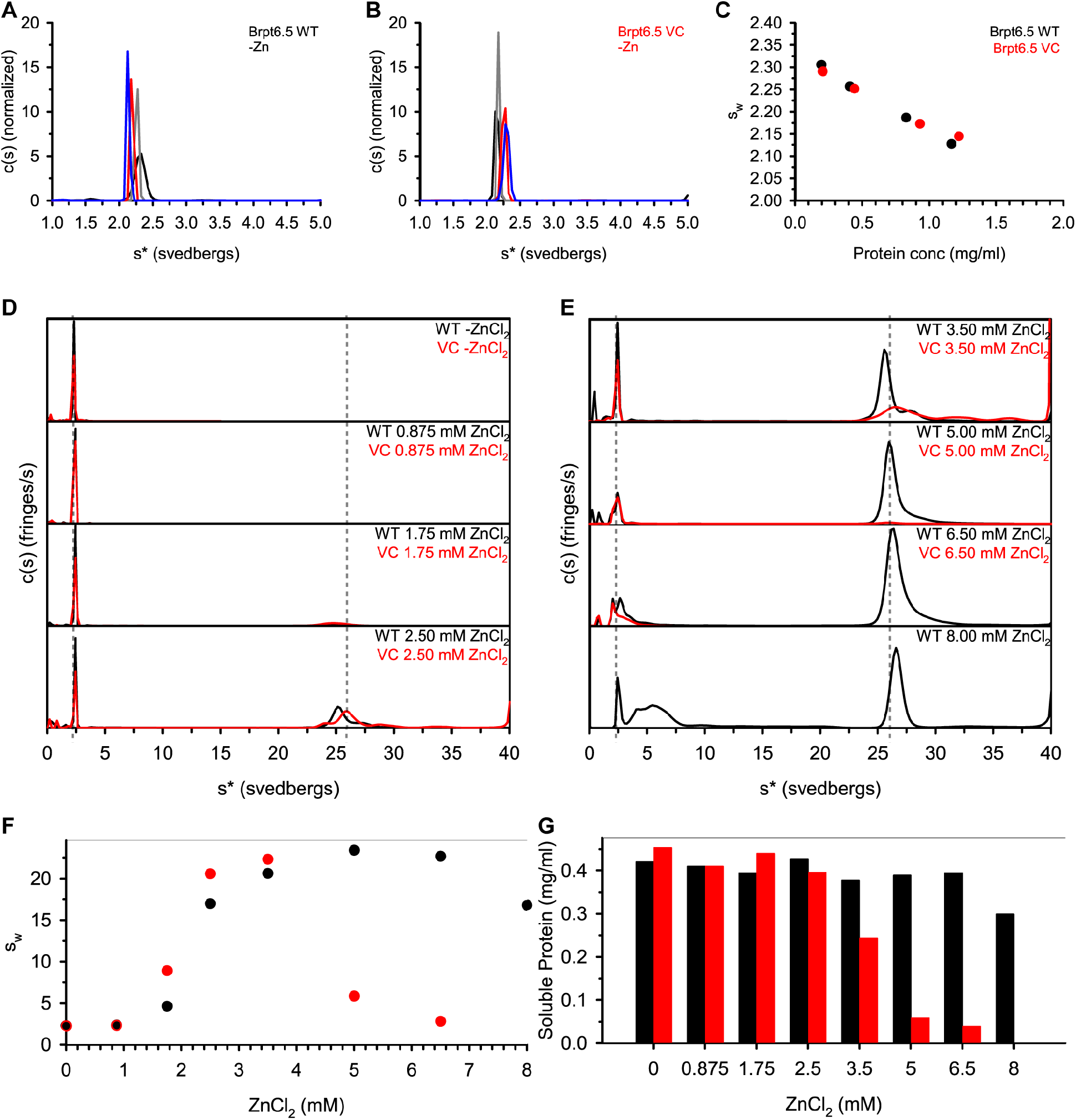
The C-terminal half-repeat cassette is not responsible for the formation of the large ~26 S species. A range of concentrations of Brpt6.5 WT (A) and Brpt6.5 VC (B) were examined in the absence of ZnCl_2_. The resulting sw values are plotted against protein concentration in panel (C). Brpt6.5 WT and VC mutant were then examined over a wide range of ZnCl_2_ concentrations. (D) and (E) show the c(s) distributions, with WT in black and VC in red. At 8.00 mM ZnCl_2_, VC was too aggregated after dialysis into the buffer to perform the experiment. Dashed lines are shown at s* = 2.2 (monomer) and s* = 26 for reference. (F) The sw obtained from c(s) distributions in (D) and (E) plotted against ZnCl_2_ concentration. (G) A bar chart shows the protein concentration in each experiment determined by integration of the area under the c(s) distribution from approximately 1 s* to 38 s*. For (F) and (G), the WT data are shown in black, while the VC data are shown in red.

The ~26 S species were unexpected based on what has been previously observed for Brpt5.5 from strain RP62A Aap – which involved a monomer (~2 S), dimer (~4 S), and tetramer (~7 S) before aggregating into fibrils which sediment essentially immediately in the AUC. A His6-MBP-tagged Brpt3.5 did show highly elongated and compact species sedimenting beyond ~7 S, however, this was a highly heterogeneous mixture of species that required incubation with ZnCl_2_ at 37 °C (21). Like the Brpt5.5 dimer and tetramer, the Brpt6.5 ~26 S species are reversible upon addition of DTPA (diethylenetriaminepentaacetic acid) to chelate the Zn^2+^ (Figure S3) (21). Due to the previously defined importance of the C-terminal G5 subdomain on assembly and stability (7), we hypothesized that the variant sequence in the C-terminal G5 subdomain of Brpt6.5 from strain 1457 Aap may be influencing the assembly or energy landscape and causing the ~26 S species to form as stable intermediates, rather than allowing for direct aggregation from tetramer into fibrils as previously reported (20). To test this hypothesis, a mutant Brpt6.5 was designed where the C-terminal G5 subdomain variant sequence was swapped for the consensus sequence, resulting in a completely consensus Brpt6.5 construct (referred to as Brpt6.5 VC: **<u>V</u>**ariant sequence mutated to **<u>C</u>**onsensus sequence).

Firstly, the Brpt6.5 VC construct was evaluated by CD, which confirmed proper folding and nearly identical thermal stability as WT (Figure S4, Table S2). While previous studies using Brpt1.5 constructs showed an increase in thermal stability of 6 °C when a variant sequence is present in the C-terminal G5 subdomain (7), the sequence swap in the context of 6 and a half B-repeats may be expected to have a much smaller relative contribution considering the additional 5 B-repeats that are present. As with Brpt6.5 WT, Brpt6.5 VC showed very similar protein concentration dependence on sw (Figure 4B and Figure 4C). Brpt6.5 VC Zn^2+^-dependence was then evaluated in parallel with Brpt6.5 WT (i.e., using the same Zn^2+^-containing dialysis buffers to ensure matched conditions). Interestingly, Brpt6.5 VC more readily formed the ~26 S species with an increase in sw values up until significant protein aggregation was observed at 5 mM and 6.5 mM ZnCl_2_ (Figure 4). At 8 mM ZnCl_2_, no soluble protein could be recovered after dialysis. These results are consistent with previous findings for Brpt1.5, where a consensus sequence in the C-terminal G5 subdomain allowed for better Zn^2+^-dependent assembly compared to constructs with the variant sequence (7). Based on these Brpt6.5 WT and VC results in comparison to Brpt5.5 data, we conclude that the presence of an additional B-repeat, rather than the presence of a C-terminal variant repeat, is likely the reason for the presence of the stable intermediates near 26 S.

### Characterization of the large Zn^2+^-induced species of Brpt6.5

To obtain more information about the identity of the ~26 S species, sedimentation equilibrium AUC experiments were conducted. While sedimentation velocity experiments can provide accurate molecular weight estimates in many cases, there are possible complications when samples contain multiple species – especially when the species are interacting. A sedimentation equilibrium experiment was performed at 4,000 rpm, such that the ~26 S species will be able to form a concentration gradient rather than sediment completely. At such a low speed, the monomer is not expected to form an appreciable gradient above the level of noise. More importantly, equilibrium experiments determine molecular weight of species directly, without needing to estimate the shape or diffusion coefficient of the species, as is the case for sedimentation velocity.

Such an equilibrium experiment examining Brpt6.5 WT at three loading concentrations between 0.5 mg/ml and 0.1 mg/ml with 3.50 mM ZnCl_2_ at 4,000 rpm is shown in Figure S5. Because it is unclear whether the reaction boundaries sedimenting near 26 S in the sedimentation velocity experiments represent just two species or if there are multiple oligomers or conformations, a single-species fit was used to determine the weight-average molecular weight of the 26 S oligomer(s). Based on the sedimentation velocity distributions obtained under similar conditions (Figure 3 and Figure 4), the sample is expected to contain monomer and the larger species sedimenting near 26 S. Because the monomer is not expected to contribute significantly to the concentration gradient (due to the low buoyant MW), the observed gradient would only be dependent on the larger oligomers present in the mixture. The best fit to a single ideal species model indicated a weight-averaged MW of 2.61 MDa (2.53-2.70 MDa). As expected, inclusion of a non-interacting monomer in the model did not impact the fit (Table S3). A weight-average molecular weight of 2.61 MDa lies between a 28-mer (2.57 MDa) and a 32-mer (2.93 MDa), if we assume the tetramer is the building block, as described in further detail below. Therefore, one possible interpretation of the equilibrium data is that this sample contains a mixture of 28- and 32-mers. The sedimentation coefficients of the two reaction boundaries (25 S and 27 S) observed in the SEDFIT c(s) analysis of the sedimentation velocity data may therefore represent the 28-mer and 32-mer. Fitting only the faster-sedimenting species in the 32,000 rpm experiment (Figure 3B, Figure S6) provided a weight-average frictional ratio of 2.67. Assuming this frictional ratio for the 28-mer and 32-mer, the two species would sediment at 25 S and 27 S – consistent with the observed reaction boundaries (See Methods and Table S4).

The hydrodynamic radius (*R_h_*) of the ~26 S species was measured by DLS. A sample of Brpt6.5 in the presence of 3.50 mM ZnCl_2_ was examined at 20 °C (Figure S7A and Figure S7B). While sedimentation velocity AUC was able to separate the monomer-dimer-tetramer species from the larger oligomer, DLS was unable to resolve the species. This resulted in the DLS volume and intensity distributions showing a single peak, which was composed of all oligomers present. The resulting Z-average *R_h_* at 20 °C was 27 nm. At 38 °C, the distribution shifted, while still showing as a single peak with a polydispersity index just above 0.1, and an *R_h_* of 35 nm. As temperature increases, the population of the larger oligomer is expected to increase, resulting in a slightly higher *R_h_* value. Interestingly, at 40 °C, a major increase in the polydispersity index was observed (Table S5), with distributions showing high heterogeneity indicative of aggregation (Figure S7). Interestingly, Brpt5.5 also aggregates in the same temperature range when present with the same ZnCl_2_ concentration (20).

### The mechanism of Brpt6.5 dimerization

The dimerization of B-repeats from strain RP62A Aap has been very well-characterized (7,13,18–20,37). Biophysical data on Brpt1.5 constructs first determined the presence of a histidine in the Zn^2+^-binding site, then showed that 1-2 Zn^2+^ ions are bound per G5 subdomain during dimerization (13,37). X-ray crystallography structures then provided high-resolution detail of the Zn^2+^-binding site, and also confirmed that B-repeats occupy a highly extended global conformation as monomers and form a dimer in a side-by-side orientation with essentially no change in the conformation of the protein (7,18). Brpt5.5 was found to have a similar dimerization mechanism as Brpt1.5 and Brpt2.5 constructs, with 1-2 Zn^2+^ ions binding per G5 subdomain during dimerization. Biophysical and SAXS analyses of the Brpt5.5 monomer and dimer confirmed a highly extended monomer and a similar side-by-side dimer (19,20,22).

To understand the dimerization of Brpt6.5 from strain 1457 Aap, a construct was designed which would be unable to dimerize if the mechanism of dimerization is consistent with RP62A Aap B-repeats. This construct, referred to as Brpt6.5 7xE203A, contained the E203A substitution that can completely inhibit Brpt1.5 dimerization (18). Despite pleiomorphic coordination of Zn^2+^ observed in multiple crystal structures of the Brpt1.5 dimer, E203 directly ligated Zn^2+^ in each structure. The Brpt6.5 7xE203A construct not only contains the E203A substitution consistent with the Brpt1.5 mutation previously examined, but also contains alanine substitutions in the equivalent position of each of the 6 additional G5 sub-domains of Brpt6.5.

The Brp6.5 7xE203A mutant was monomeric in the absence of Zn^2+^, as expected (Figure 5A, Figure 5D). It was unable to dimerize in the presence of Zn^2+^ (Figure 5B, Figure 5E), with only a minor increase in the sedimentation coefficient that has consistently been observed previously and is likely the result of Zn^2+^ binding to the monomer (18,19). This is clearly not indicative of dimerization, as there is no significant increase in sw with protein concentration in the presence of Zn^2+^ (Figure 5C, Figure 5D) or with increasing Zn^2+^ concentrations (Figure 5B, Figure 5E). Given the identical ability of the H75A/E203A mutations to inhibit Zn^2+^-dependent dimerization, it is concluded that B-repeats from strain 1457 Aap undergo dimerization via a mechanism similar to B-repeats from strain RP62A Aap.

**Figure 5.**
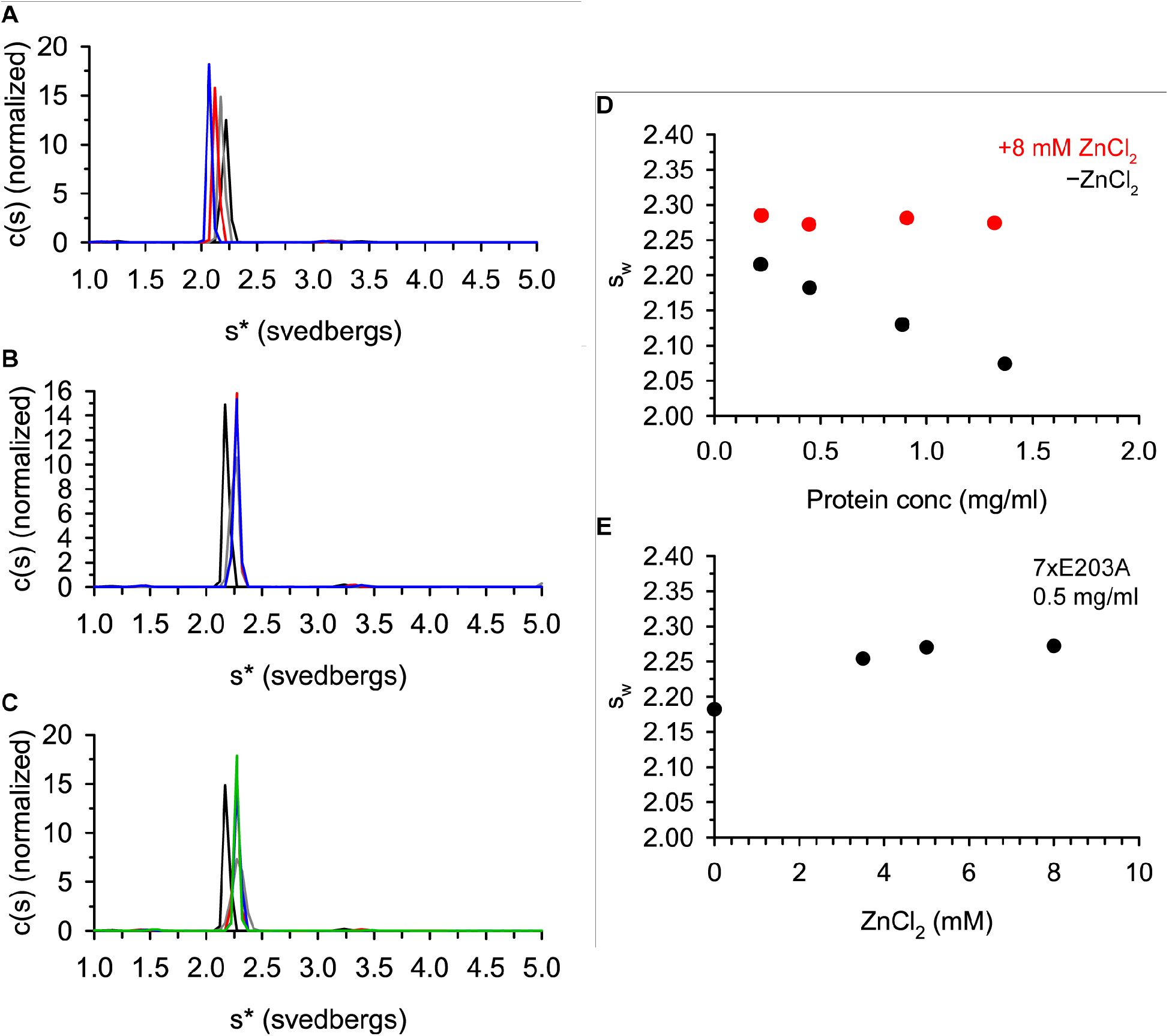
Brpt6.5 7xE203A is unable to dimerize. (A) Brpt6.5 7xE203A analyzed in the absence of Zn^2+^ at increasing protein concentrations (0.22 mg/ml = black line, 0.45 mg/ml = grey line, 0.89 mg/ml = red line, 1.37 mg/ml = blue line). (B) The c(s) distributions from samples of 0.5 mg/ml Brpt6.5 7xE203A at increasing ZnCl_2_ concentrations (-ZnCl_2_ = black line, 3.50 mM ZnCl_2_ = grey line, 5.00 mM ZnCl_2_ = red line, 8.00 mM ZnCl_2_ = blue line). (C) Distributions shown are 7xE203A at increasing protein concentrations in the presence of 8 mM ZnCl_2_ (0.22 mg/ml = grey line, 0.45 mg/ml = red line, 0.91 mg/ml = blue line, 1.32 mg/ml = green line). 7xE203A (0.45 mg/ml) without ZnCl_2_ (black line) is shown for reference. Distributions for samples in 8 mM ZnCl_2_ overlay very closely, while the -Zn distribution is shifted to a lower s-value. The sw value from the distributions in (A) – (C) are shown in (D) and (E). (D) shows the sw relationship to protein concentration in the absence of ZnCl_2_ and in the presence of 8.00 mM ZnCl_2_. (E) shows sw versus ZnCl_2_ concentration, while maintaining a constant 7xE203A concentration.

### H85-dependency of tetramer formation

Recent reports have shown the critical importance of the Brpt5.5 tetramer for the ability of Aap to form functional amyloid fibrils (20,21). The Brpt5.5 tetramer is formed by two dimers coming together in a side-by-side fashion, and tetramerization is mediated via Zn^2+^-binding to a histidine in position 85 of the spacer subdomain of each B-repeat. These conclusions are based on rigorous characterization of single- and multi-H85A mutants via biophysical and SAXS analyses (20,22). It was noteworthy that Brpt1.5 and Brpt2.5 are not observed in the tetramer state, or as aggregate, indicating that an important factor in tetramer assembly is the number of B-repeats present in the construct.

To understand tetramer assembly of Brpt6.5, a mutant was constructed containing the H85A mutation in all six spacer subdomains, called 6xH85A. Note that the C-terminal half-repeat consists of only a G5 subdomain, leaving only six spacer subdomains (and therefore six H85 positions) in Brpt6.5. This construct is analogous to the Brpt5.5 5xH85A construct, which was limited in assembly to the dimer state by preventing Zn^2+^-coordination *in trans* via the H85 residues (20,22). The Brpt6.5 6xH85A mutant is monomeric in the absence of Zn^2+^ (Figure 6A, Figure 6B). The Zn^2+^-dependent assembly of Brpt6.5 6xH85A was directly compared with the previously characterized Brpt5.5 5xH85A (Figure 6C and Figure 6D). There is a clear difference in the sw values between the two proteins, with Brpt5.5 reaching nearly complete dimer at 3.5 mM ZnCl_2_, while Brpt6.5 shows only a small fraction of assembly. This major weakening of the dimer assembly observed for Brpt6.5 6xH85A compared to Brpt5.5 may be an effect of the variant C-terminal G5 subdomain, given that Brpt1.5 constructs with a C-terminal variant sequence exhibited extremely weak dimerization (7).

**Figure 6.**
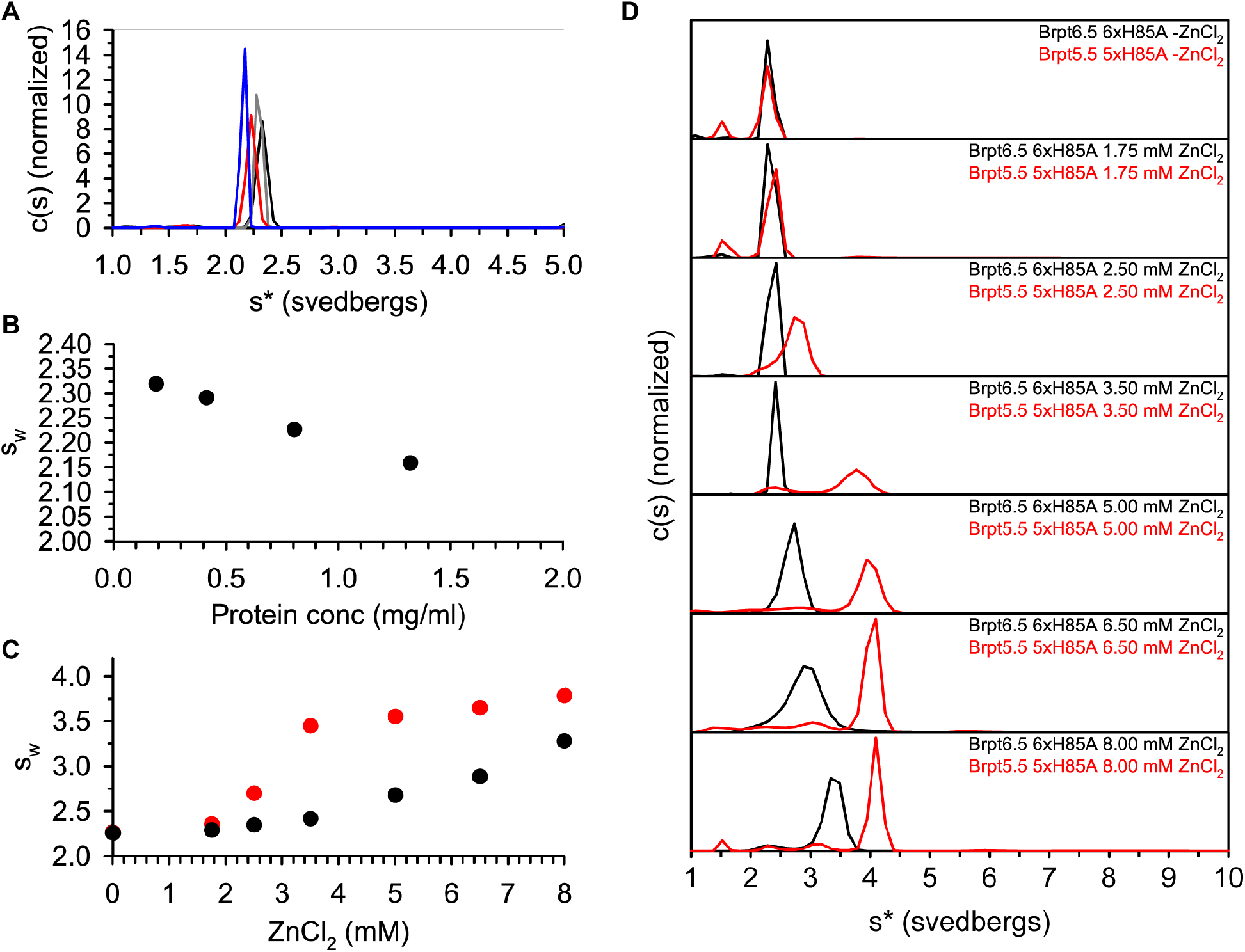
Brpt6.5 6xH85A is unable to form the tetramer and larger species. Panel (A) shows c(s) distributions of Brpt6.5 6xH85A in the absence of Zn^2+^ at increasing protein concentrations (0.19 mg/ml = black line, 0.41 mg/ml = grey line, 0.80 mg/ml = red line, 1.32 mg/ml = blue line). (B) Shows the sw plotted against protein concentration. (C) The sw of Brpt6.5 6xH85A or Brpt5.5 5xH85A (strain RP62A) at multiple ZnCl_2_ concentrations determined from c(s) distributions shown in (D). (D) The c(s) distributions for Brpt6.5 6xH85A are shown in black, while Brpt5.5 5xH85A are shown in red. AUC experiments were performed in 50 mM MOPS pH 7.2, 50 mM NaCl. Where ZnCl_2_ is present, it was included in the dialysis buffer and the protein concentration was approximately 0.5 mg/ml.

### Periodicity of tetramer formation in single-H85A mutants

Recent work on Brpt5.5 from strain RP62A revealed that single-H85A mutants in every other B-repeat spacer subdomain could completely inhibit tetramer formation (22). This was consistent with the proposed tetramer structure being formed by two dimers lining up side-by-side, each of which are twisting down the axis (20,22). This particular tetramer orientation places alternating H85 positions along the length of the construct at the interface of the two dimers, where Zn^2+^ ions can be coordinated across dimers to form the tetramer. The remaining H85 positions protrude away from the tetramer and can mediate downstream aggregation into fibrils (22).

In order to determine whether or not Brpt6.5 utilizes a similar tetramer-dependent assembly process, single-H85A mutants were constructed and their assembly examined by sedimentation velocity AUC at several ZnCl_2_ concentrations (Figure 7). We observe that appropriate H85A mutants of Brpt6.5 can form a tetramer species like Brpt5.5 if assembly to the larger ~26 S species is curtailed. Overall, the assembly data reveal a periodicity similar to that seen for to Brpt5.5 (22), where, for example, H213A supports tetramer, but H341A cannot form tetramer. In addition, a H85A mutation in either the N-terminal spacer subdomain or the C-terminal spacer subdomain is insufficient to prevent formation of the ~26 S species (Figure 7A and Figure 7F). This could mean that these H85 positions are not in the interface of the ~26 S species, or that contrary to Brpt5.5, the extra B-repeat of Brpt6.5 reduces the importance of a single H85A mutation on overall assembly. No signal was observed between dimer (~4 S) and tetramer (~7 S) for H85A, H341A, and H597A. The other single-H85A mutants showed formation of tetramer, with reaction boundaries in the ~4 S to ~7 S range. Therefore, while the presence of the ~26 S species complicates this analysis slightly, the same periodicity is observed for Brpt6.5 – a convincing indication that the tetramer is formed in a manner similar to Brpt5.5, but that Brpt6.5 tetramers further assemble into the ~26 S species with such strong cooperativity that isolated WT Brpt6.5 tetramer is not observed. These data also highlight the importance of the middle four H85 positions in formation of the ~26 S species, whether they were involved in tetramer formation or not.

**Figure 7.**
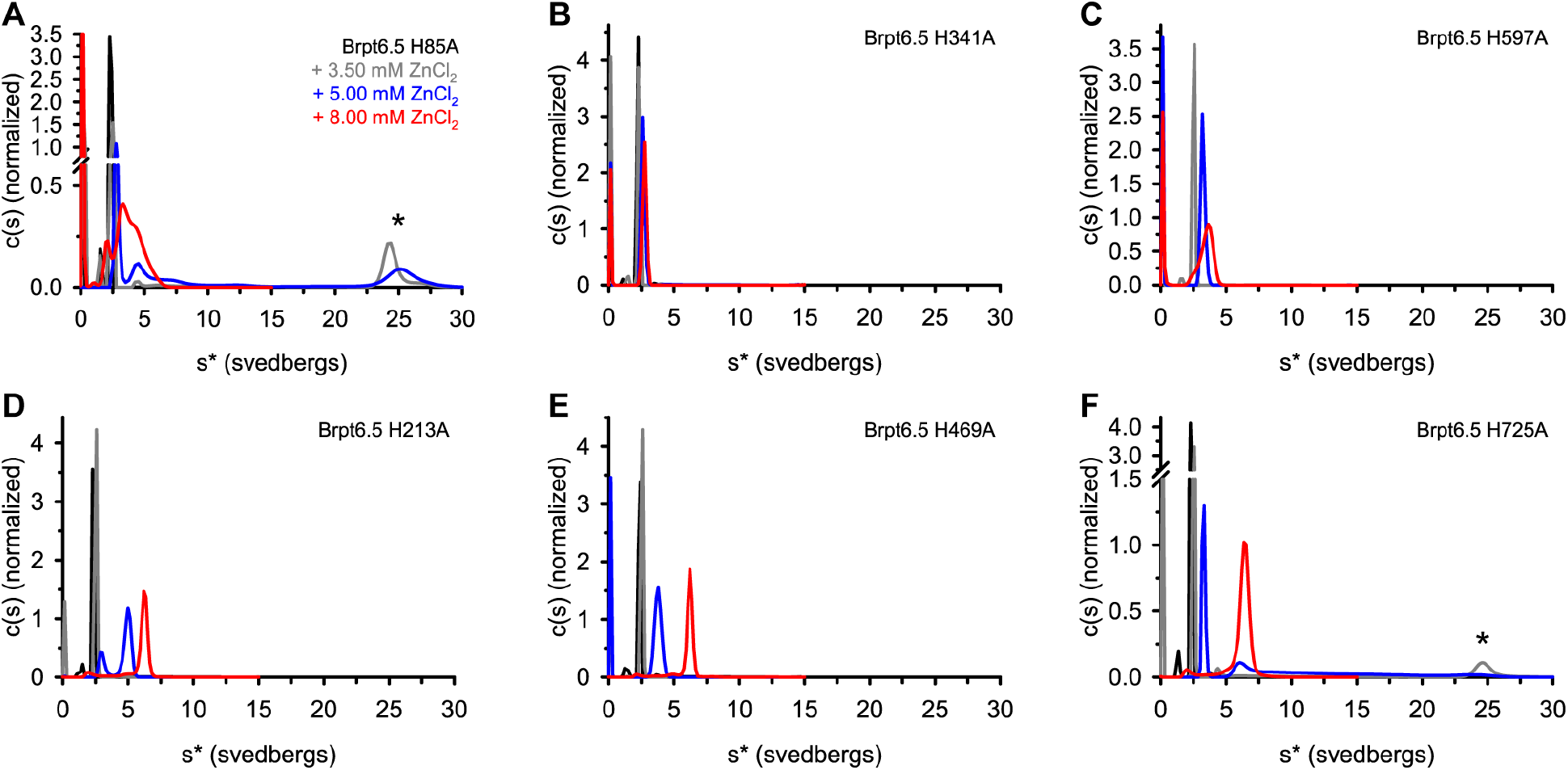
Brpt6.5 single-H85A mutants reveal which positions support tetramer formation. The c(s) distribution obtained from sedimentation velocity AUC experiments are shown for each of the single-H85A mutants that compose the 6xH85A mutant. Experiments were performed in the absence of Zn^2+^ (black line) or in the presence of 3.50 mM ZnCl_2_ (grey line), 5.00 mM ZnCl_2_ (blue line), and 8.00 mM ZnCl_2_ (red line). Data were collected in 50 mM MOPS pH 7.2, 50 mM NaCl at approximately 0.5 mg/ml protein. ZnCl_2_ was dialyzed into the sample when present. Brpt6.5 H85A (A) and H725A (F) show the presence of the large ~26 S species (marked with an asterisk (*)). A single H85A mutation in the third (B – H341A) or fifth (C – H597A) spacer subdomain prevent formation of the tetramer (~7 S) under these conditions. In contrast, a single H85A mutation in the second (D – H213A) or fourth (E – H469A) spacer subdomain allow for formation of the tetramer but prevent formation of the ~26 S species.

### Brpt6.5 aggregation and amyloidogenesis

A primary function of Aap B-repeats is to assemble and aggregate into amyloid fibrils. Characterization of Brpt3.5 and Brpt5.5 aggregation has been reported using several approaches (20,21). Presented in Figure 8A is a turbidity assay, wherein small aliquots of ZnCl_2_ are titrated into a sample of Brpt6.5 WT, 7xE203A, 6xH85A, or buffer, and the turbidity at 400 nm is reported after each addition. As Zn^2+^-induced aggregation occurs, turbidity increases. Brpt6.5 WT readily aggregates beyond ~7-10 mM Zn^2+^, whereas the Brpt6.5 6xH85A requires much higher Zn^2+^ concentrations. These results are consistent with Brpt5.5 results previously published (20,22). Interestingly, Brpt6.5 7xE203A exhibits aggregation with a sigmoidal trend at intermediate ZnCl_2_ concentrations, despite its inability to dimerize. This is a critical observation that could suggest that B-repeats may bypass the E203A-mediated dimer when at higher Zn^2+^ concentrations, allowing aggregation to occur via H85-mediated assembly. It could be the case that the presence of H85 in each B-repeat is still able to mediate downstream aggregation in a Zn^2+^-dependent manner.

**Figure 8.**
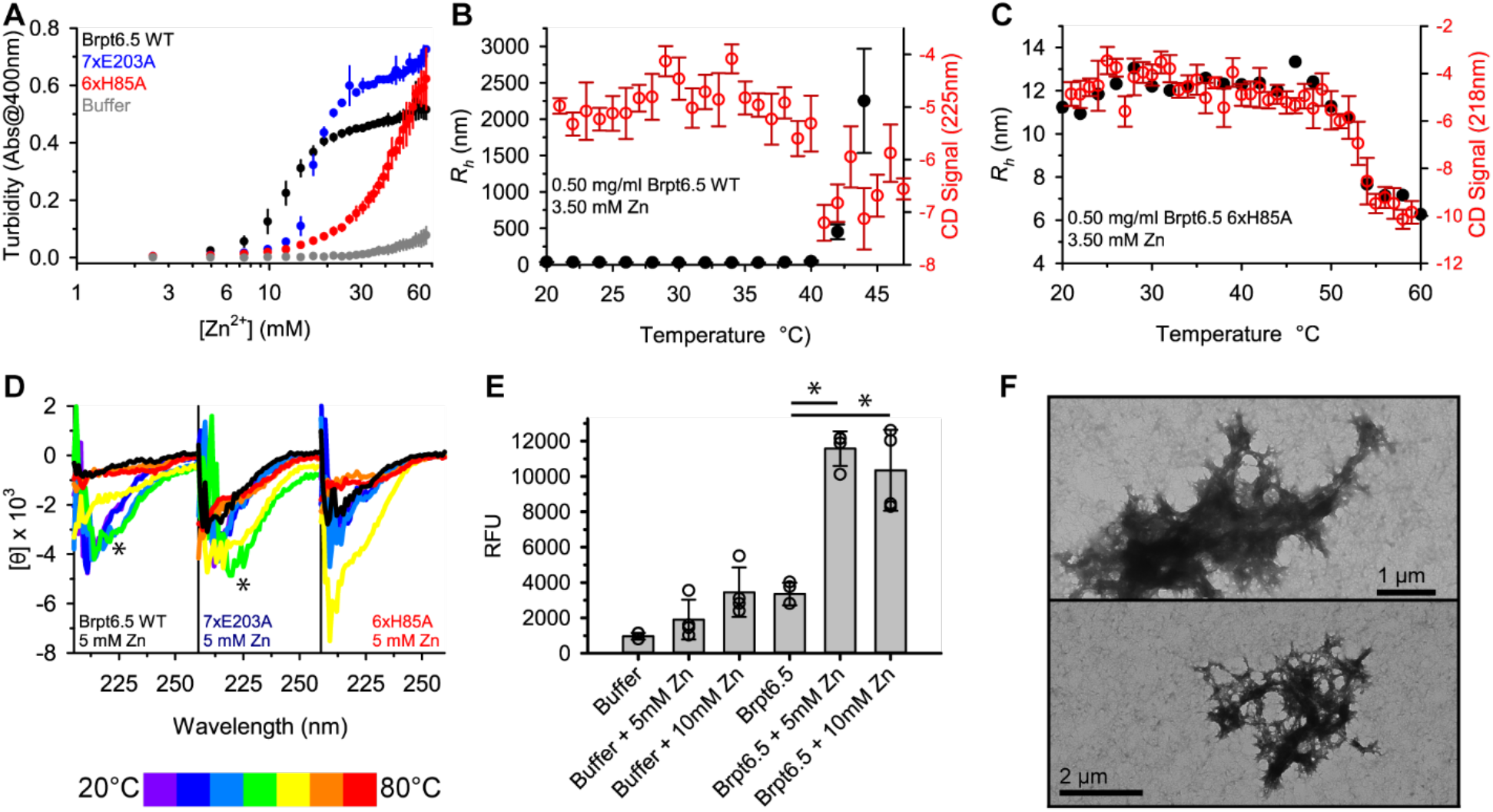
Brpt6.5 from strain 1457 aggregates into amyloid-like fibrils using a similar mechanism as Brpt5.5 from Aap from strain RP62A. (A) The light scattering of each Brpt6.5 construct in the presence of increasing ZnCl_2_. Triplicate measurements were performed, and the error bars show ± 1 standard deviation (SD). Temperature-dependent aggregation of Brpt6.5 WT (B) and Brpt6.5 6xH85A (C) in the presence of Zn^2+^ was measured by DLS and the data presented as radius of hydration (*R_h_*). The DLS data are presented as black markers (average of three measurements) with error bars showing ± 1 SD. (B) and (C) also show the CD signal in machine units as red markers with error bars showing the error reported by the instrument. The CD wavelength monitored in (B) was 225 nm, while 218 nm was monitored in (C). (D) shows Far-UV CD measurements for each construct in the presence of 5 mM ZnCl_2_ in 10 °C increments from 20 °C - 80 °C. Asterisks mark the presence of a 225 nm minimum observed for Brpt6.5 WT (40 °C) and 7xE203A (50 °C). The black line shows a scan taken at 20 °C after heating to 80 °C and indicates the degree to which aggregation was reversible. (E) Fluorescence of Thioflavin T (ThT) measured for samples after 24 hours at 37 °C with shaking. The bar chart indicates the average of 4 replicates, with error bars showing ± 1 SD. The markers indicate the 4 individual data points for each sample. Asterisks denote significant differences between Brpt6.5 WT without Zn^2+^ and either in the presence of 5 mM Zn^2+^ or 10 mM Zn^2+^ according to a two-tailed, two-sample unequal variance Student’s *t* test with *p* < 0.05. (F) TEM micrographs of Brpt6.5 + 10 mM ZnCl_2_ from the ThT assay shown in (E).

A second approach to characterizing the aggregation propensity of Brpt6.5 is to start with Zn^2+^ present in the sample, then use DLS to follow aggregation in a temperature-dependent fashion. In parallel, the same sample can be monitored by CD to follow the formation of a minimum at 225 nm that has been associated with the formation of amyloid fibrils (20,21,38,39). For Brpt6.5 WT (Figure 8B) both DLS-measured aggregation and the appearance of a minimum in the CD signal at 225 nm occur concomitantly at ~42 °C. On the contrary, Brpt6.5 6xH85A shows only unfolding from dimer and tetramer species to random coil by DLS, along with development of a minimum measured at 218 nm near 52 °C (Figure 8C), consistent with unfolding observed in the absence of Zn^2+^ (Figure 2). Figure 8D shows Far-UV CD wavelength scans conducted on similar samples as the DLS/CD experiments. The temperature was increased in 10 °C increments, revealing the appearance of the 225 nm minimum for Brpt6.5 WT and Brpt6.5 7xE203A, whereas Brpt6.5 6xH85A simply unfolded. In agreement with the Zn^2+^ turbidity assay that showed 7xE203A required slightly higher Zn^2+^ concentrations than WT, the 7xE203A required slightly higher temperatures to induce the fibril-related conformational change. The DLS/CD observations are nearly identical to those observed for Brpt5.5, suggesting a very similar H85-mediated aggregation mechanism despite the different distributions of oligomeric states between the two B-repeat constructs (20–22).

Lastly, the aggregate formed by Brpt6.5 was examined in comparison to that previously published for Brpt5.5 (21). A fluorescent amyloid-detecting dye, Thioflavin T (ThT), was used in conjunction with transmission electron microscopy (TEM) to show the formation of amyloid-like fibrils. Figure 8E demonstrates that a significant increase in ThT fluorescence is observed for Brpt6.5 WT in the presence of Zn^2+^ after a 24-hour incubation at 37 °C. Examination of the ThT-positive sample of Brpt6.5 WT with 10 mM ZnCl_2_ by TEM (Figure 8F) showed aggregates similar in morphology to those observed for Brpt5.5 and in native *S. epidermidis* RP62A biofilms (21).

## DISCUSSION

This manuscript describes an investigation of the mechanism of assembly and amyloidogenesis of the complete Aap B-repeat superdomain from *S. epidermidis* strain 1457. This is the first report to use a protein construct containing the complete B-repeat superdomain of Aap or SasG (the *S. aureus* ortholog) for biophysical analyses. While the mechanisms of B-repeat assembly and amyloidogenesis have been published previously, all these studies were performed using protein constructs based on B-repeats from Aap from *S. epidermidis* strain RP62A (7,13,18–21,37). Most of the biological experiments designed to understand the role of Aap have used *S. epidermidis* strain 1457, which is amenable to genetic manipulation (6,8,9,11,23,40). Therefore, it is important to understand if there are differences in B-repeat assembly and amyloidogenesis across strains, so that the effects of Aap or B-repeat mutations can be evaluated reliably between biophysical, structural, and functional experiments. This work fills that gap in knowledge.

The evaluation presented in this study indicates that B-repeats from Aap from strains RP62A and 1457 behave similarly in terms of overall Zn^2+^-induced assembly into amyloid-like fibrils. There are some very intriguing differences, however. One such difference is the concentration of Zn^2+^ required for dimerization between Brpt6.5 6xH85A and Brpt5.5 5xH85A. Based on previous work, the variant sequence in the C-terminal G5 subdomain was shown to be critical for dimerization (7), but that was in the context of a Brpt1.5 construct. Clearly, the impact is not as significant in the context of Brpt6.5. Also factoring into dimerization is the number of B-repeats – where constructs containing a higher number of B-repeats are expected to dimerize in the presence of lower Zn^2+^ concentration due to the chelate effect (19,41). The most obvious difference is the ability for Brpt6.5 to form very large, stable and reversible 28-mer and 32-mers that can then undergo irreversible assembly into amyloid fibrils as with Brpt5.5. On the other hand, Brpt5.5 has a terminal stable species of tetramer, which then undergoes aggregation into fibrils without any detectable larger stable intermediates (20). If Brpt5.5 can in fact form similar higher-order oligomers like Brpt6.5, these species must be very transient such that they are rapidly pushed toward amyloid fibril assembly. In contrast, experiments using a series of H85A-containing mutant constructs showed that Brpt6.5 also assembles through a tetramer species, but it assembles further with strong cooperativity to the 28-mer and 32-mer species, such that no free Brpt6.5 tetramer is observed. A possible interpretation of the formation of these stable species is that they form as a branching or ring-like structure, such that after 28 or 32 molecules are assembled, there is limited space for additional tetramers to be added. At that point, additional energy is required to push the system further toward amyloid fibril formation.

Despite these intriguing differences between in the energetics of B-repeat Zn^2+^-dependent assembly, both the Brpt5.5 and Brpt6.5 constructs aggregate into amyloid fibrils under very similar Zn^2+^ concentrations and temperatures (~40 °C). This is a very important consideration for the biological implications of B-repeat assembly. Although different strains of *S. epidermidis* express Aap containing a wide variety of the number and cassette identity of B-repeats, there appear to be natural mechanisms in place to control the energy landscape of assembly and balance out the ability of Aap to mediate biofilm formation only under the desired environmental conditions. While the enzymatic cleavage of the A-repeats and lectin domain is a major switch for Aap to mediate attachment or biofilm accumulation, there is clearly also a potential for control via the B-repeat superdomain itself.

Implications of this work are that B-repeats from Aap across *S. epidermidis* strains may differ in the distribution of oligomeric states formed in the presence of Zn^2+^, but that they share the common characteristic of nucleating functional amyloid fibers, which is not surprising based on their high sequence identity and the presumed biological importance of functional amyloid for biofilm mechanical strength. Therefore, for genetic manipulation of *S. epidermidis* 1457, the findings from B-repeats from Aap from strain RP62A should be translatable. For example, to test the impact of Aap amyloidogenesis on biofilm formation, one could produce H85A mutations in each B-repeat of Aap. A very important observation from this work is that while the E203A mutations prevented dimerization, aggregation was still observed at intermediate Zn^2+^ concentrations. This means that to test the impact of dimerization on biofilm formation, a genetically modified *S. epidermidis* 1457 strain should contain both the E203A and H85A mutations in the Aap B-repeats. While the *S. aureus* SasG B-repeats have only been biophysically and structurally analyzed in the monomer state, this work provides a guide to anticipating what variables may be important to consider for Zn^2+^-mediated assembly and biofilm formation in B-repeat surface proteins found in other staphylococcal species.

## Acknowledgements

The authors thank Dr. Catherine Shelton and Dr. John Burgner for offering comments on the manuscript. We also thank Dr. Peter Sherwood and Dr. Walter Stafford for implementing updates within SEDANAL that allow for loading and analyzing greater than 999 scans. This is particularly useful when using interference optics to collect data using an equilibrium method without any time delay between scans to perform velocity experiments (99 steps of 99 scans).

