## Supplemental Data for "The assembly landscape of the complete B-repeat superdomain from *Staphylococcus epidermidis* strain 1457"

[#] Current affiliation: BioAnalysis, LLC, Philadelphia, PA 19134, USA

### Supporting Material

Figures:
Figure S1. Summary of B-repeat constructs used in this study.
Figure S2. Sequence alignment of regions of Aap from *S. epidermidis* strain RP62A and 1457.
Figure S3. The large oligomers sedimenting near 26 S are reversible upon $Zn^{2+}$-chelation.
Figure S4. Swapping the sequence cassette from variant to consensus does not affect secondary structure or stability.
Figure S5. AUC-EQ analysis of Brpt6.5 WT + $ZnCl_2$.
Figure S6. Analysis of Brpt6.5 WT + 3.50 mM $ZnCl_2$ at 32,000 rpm.
Figure S7. DLS distributions of Brpt6.5 WT + $ZnCl_2$ at increasing temperature.

Tables:
Table S1. AUC hydrodynamic nonideality results.
Table S2. CD thermal denaturation results.
Table S3. AUC-EQ fit statistics.
Table S4. Predicted Sedimentation Coefficients for oligomers of Brpt6.5.
Table S5. DLS measurements of particle size.

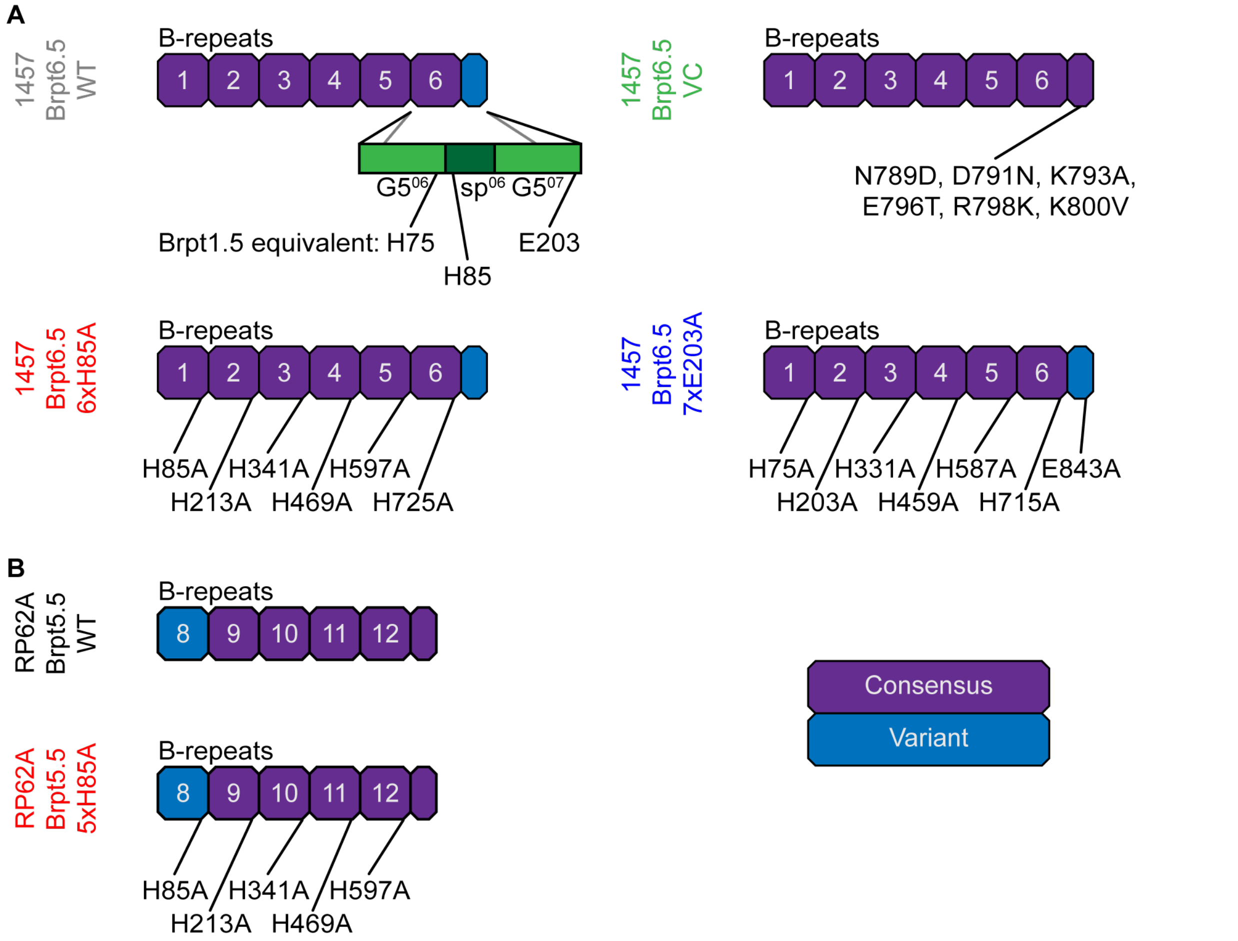


**Figure S1. Summary of B-repeat constructs used in this study.** (A) Brpt6.5 WT and mutants from strain 1457. (B) Brpt5.5 WT and the 5xH85A mutant. Note that the Brpt5.5 WT construct was not used in this study and is shown only for reference. See Yarawsky & Herr (1) for more information on Brpt5.5 5xH85A. B-repeats are colored according to the consensus or variant subtype, as described in the text and in Shelton, et al. (2).

**Table S1. AUC hydrodynamic nonideality results.**

| Sample | $s^0$ | $k_s$ | rmsd |
|---|---|---|---|
| WT | 2.341 ± 0.004 | 86.2 ± 2.7 | 0.0030 |
| VC | 2.322 ± 0.007 | 69.7 ± 4.1 | 0.0046 |
| 6xH85A | 2.352 ± 0.004 | 68.0 ± 2.2 | 0.0029 |
| 7xE203A | 2.242 ± 0.002 | 59.4 ± 1.0 | 0.0012 |

Determined in 50 mM MOPS pH 7.2, 50 mM NaCl at 20 °C. Analysis performed using SEDNTERP v3.0.3 (3).

```
A-repeats
RP62A      AEEGSNAEAPQSEPTKAEEGGNAEAAQSEPTKAEEGGNAEAPQSEPTKAEEGGNAEAAQS   60
1457       AEEGGNAEAPQSEPTKAEEGGNAEAPQSEPTKTEEGSNVKAAQSEPTKAEEGSNAEAPQS   60
           ****.******************** ******:***.*.:* **********.**** **

RP62A      EPTKTEEGSNVKAAQSEPTKAEEGSNAEAPQSEPTKTEEGSNAKAAQSEPTKAEEGGNAE   120
1457       EPTKTEEGSNAKAAQSEPTKAEEGDNAEAPQSEPTKTEEGSNAKAAQSEPTKAEEGGNAE   120
           **********.*************.***********************************

RP62A      AAQSEPTKTEEGSNAEAPQSEPTKAEEGGNAEAPQSEPTKTEEGGNAEAPNVPTIK       176
1457       APQSEPTKTEEGSNAEAPQSEPTKAEEGGNAEAPQSEPTKTEEGGNAEAPNVPTIK       176
           * ******************************************************

Lectin
RP62A      IDIPPTTVKGRDNYDFYGRVDIESNPTDLNATNLTRYNYGQPPGTTTAGAVQFKNQVSFD   60
1457       IDIPPTTVKGRDNYDFYGRVDIQSNPTDLNATNLTRYNYGQPPGTTTAGAVQFKNQVSFD   60
           **********************:*************************************

RP62A      KDFDFNIRVANNRQSNTTGADGWGFMFSKKDGDDFLKNGGILREKGTPSAAGFRIDTGYY   120
1457       KDFDFNIRVANNRQSNTTGADGWGFMFSKKDGDDFLKNGGILREKGTPSAAGFRIDTGYY   120
           ************************************************************

RP62A      NNDPLDKIQKQAGQGYRGYGTFVKNDSQGNTSKVGSGTPSTDFLNYADNTTNDLDGKFHG   180
1457       NNDPLDKIQKQAGQGYRGYGTFVKNDSQGNTSKVGSGTPSTDFLNYADNTTNDLDGKFHG   180
           ************************************************************

RP62A      QKLNNVNLKYNASNQTFTATYAGKTWTATLSELGLSPTDSYNFLVTSSQYGNGNSGTYAS   240
1457       QKLNNVNLKYNASNQTFTATYAGKTWTATLSELGLSPTDSYNFLVTSSQYGNGNSGTYAD   240
           ***********************************************************.

RP62A      GVMRADLDGATLTYT                                                255
1457       GVMRADLDGATLTYT                                                255
           ***************

Pro/Gly-Rich Region
RP62A      AEPGKPAEPGKPAEPGKPAEPGTPAEPGKPAEPGTPA-----------EPGKPAEPGKPA   49
1457       AEPGKPAEPGKPAEPGKPAEPGTPAEPGKPAEPGTPAEPGTSEPGKPAEPGKPAEPGKPA   60
           *************************************           ************

RP62A      EPGKPAEPGKPAEPGTPAEPGTPAEPGKPAEPGTPAEPGKPAEPGTPAEPGKPAESGKPV   109
1457       EPGKPAEPGKPAEPGTPAEPGKPAEPGKPAEPGTPAEPGTPAEPGKPAEPGKPAEPGKPA   120
           *********************.*****************.*****.********* ***.

RP62A      EPGTPAQSGAP                                                    120
1457       EPGTPAQSGAP                                                    131
           ***********
```

**Figure S2. Sequence alignment of regions of Aap from *S. epidermidis* strain RP62A and 1457.** A sequence alignment of the A-repeats, Lectin, and Pro/Gly-Rich Region of Aap are shown. Amino acid numbering begins at zero for each region.

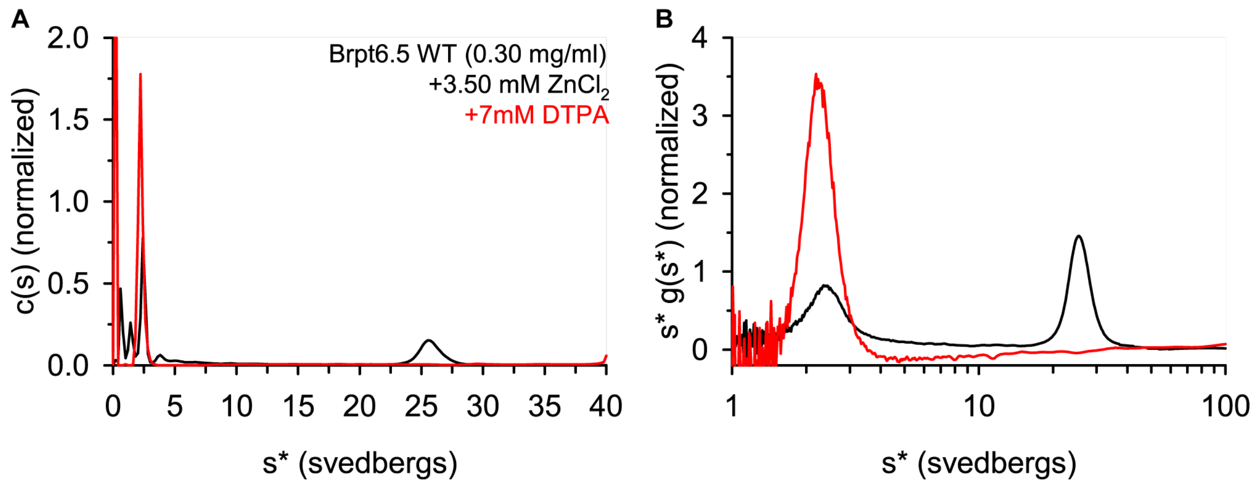


**Figure S3. The large oligomers sedimenting near 26 S are reversible upon $Zn^{2+}$-chelation.** Brpt6.5 WT was dialyzed into 50 mM MOPS pH 7.2, 50 mM NaCl, 3.50 mM $ZnCl_2$.The sample was then examined by sedimentation velocity AUC directly or with the addition of 7 mM DTPA. The final protein concentrations were approximately 0.21 mg/ml (without DTPA) and 0.12 mg/ml (with DTPA). Data were analyzed by (A) SEDFIT c(s) or (B) SEDANAL WDA.

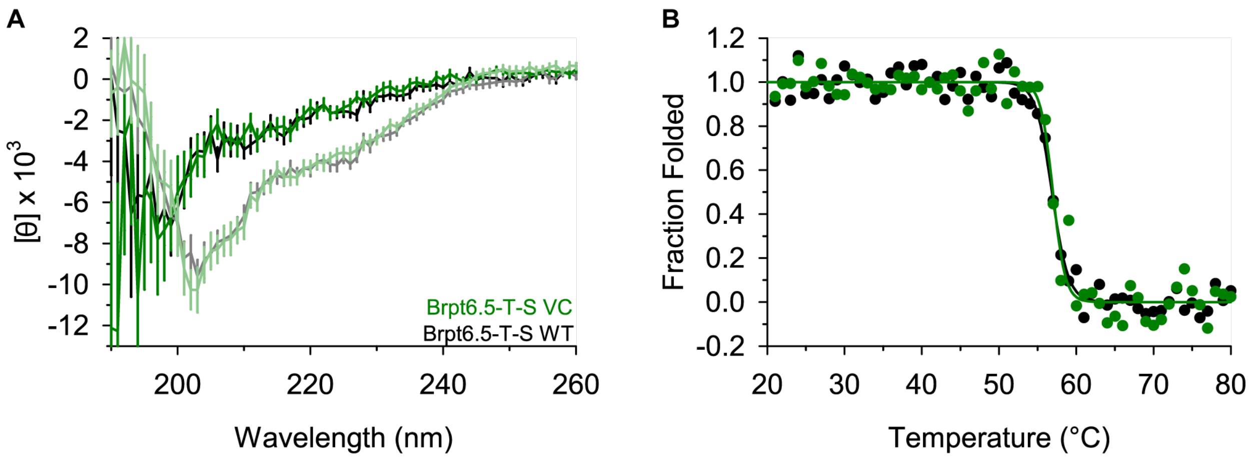


**Figure S4. Swapping the sequence cassette from variant to consensus does not affect secondary structure or stability.** (A) Far-UV CD wavelength scans showing similar secondary structure content between the WT (black line) and VC mutant (green line). Dark colored lines were obtained at 20 °C, while light colored lines were obtained at 80 °C. (B) Thermal denaturation CD experiments monitoring the CD signal at 205 nm. Each dataset was fit to a two-state model. The markers indicate data points, while the solid lines show the best fit. WT data are shown in black, while VC data are shown in green. Data are shown as fraction folded. CD data were collected in 50 mM MOPS pH 7.2, 50 mM NaCl, at 1.5 mg/ml protein.

**Table S2. CD thermal denaturation results.**

| Sample | $T_m$ (°C) | ΔH (kcal/mol) |
|---|---|---|
| WT | 56.9 ± 0.2 | -207 ± 31 |
| VC | 57.0 ± 0.2 | -272 ± 58 |

Determined at 1.5 mg/ml Brpt6.5 WT in 50 mM MOPS pH 7.2, 50 mM NaCl in a 0.1 mm cuvette.

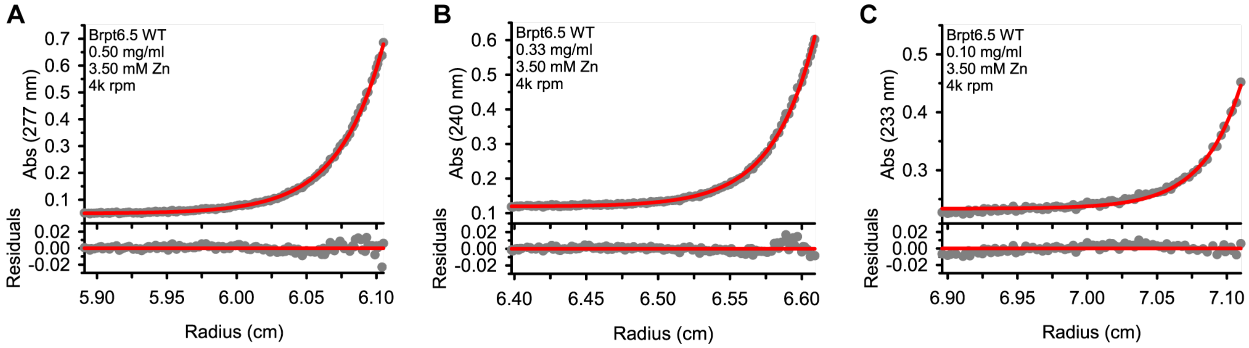


**Figure S5. AUC-EQ analysis of Brpt6.5 WT + $ZnCl_2$.** The best global fit to the single, ideal species. Panels A-C differ in the loading concentration and absorbance wavelength used for data collection. The grey markers are the raw data, while the red line is the best fit. The bottom portion of each panel shows the residuals between observed data and the fit.

**Table S3. AUC-EQ fit statistics.**

| Model | Standard Deviation | Monomer MW (kDa) | Species 1 MW (MDa) |
|---|---|---|---|
| 1 species | 3.99E-03 | - (excluded) | 2.61 (2.53 – 2.70) (fitted) |
| 1 species + monomer | 3.98E-03 | 91.72 (fixed) | 2.61 (2.53 – 2.70) (fitted) |

Determined at 0.5 mg/ml, 0.33 mg/ml, and 0.10 mg/ml Brpt6.5 WT in 50 mM MOPS pH 7.2, 50 mM NaCl, 3.50 mM $ZnCl_2$. The experiment was performed at 20 °C. Values in parenthesis indicate 95% confidence intervals reported by SEDANAL F-statistics. The monomer was incorporated as a non-interacting species without constraining the loading concentration in each dataset.

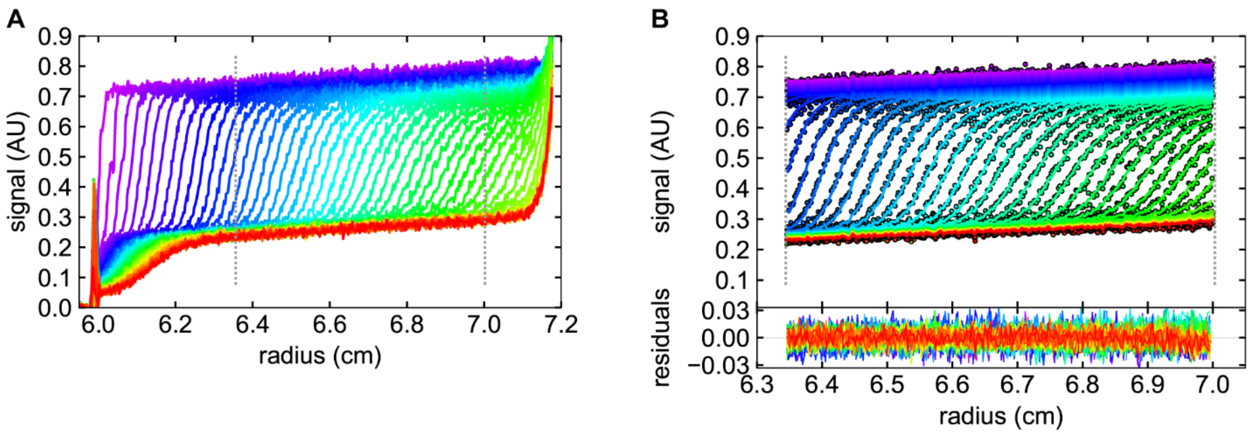


**Figure S6. Analysis of Brpt6.5 WT + 3.50 mM $ZnCl_2$ at 32,000 rpm.** The first 300 scans collected are shown in panel (A) (every $2^{nd}$ scan is shown). This particular span of scans was analyzed in order to obtain a weight-average frictional ratio for the larger species without bias from the monomer (which has an expected $f/f_0$ = 3.48 according to Table 2). Panel (B) shows the fitted region and residuals. GUSSI (4) was used to produce this figure. Every $2^{nd}$ scan is shown in each case for clarity. SEDFIT c(s) analysis was performed with TI noise fitted, but noise subtraction is turned off within GUSSI to allow better visual consistency between panels.

**Table S4. Predicted Sedimentation Coefficients for oligomers of Brpt6.5.**

| Oligomer | Sedimentation Coefficient (S) | $MW_{calc}$ (MDa) | $MW_{seq}$ (MDa) |
|---|---|---|---|
| 1 | 2.3 | 0.07 | 0.09 |
| 2 | 4.3 | 0.19 | 0.18 |
| 4 | 6.8 | 0.37 | 0.37 |
| 8 | 10.8 | 0.74 | 0.73 |
| 12 | 14.2 | 1.11 | 1.10 |
| 16 | 17.1 | 1.47 | 1.47 |
| 20 | 19.9 | 1.84 | 1.83 |
| 24 | 22.5 | 2.22 | 2.20 |
| ***28*** | ***24.9*** | ***2.58*** | ***2.57*** |
| ***32*** | ***27.2*** | ***2.95*** | ***2.93*** |
| 36 | 29.5 | 3.33 | 3.30 |
| 40 | 31.6 | 3.69 | 3.67 |
| 44 | 33.7 | 4.06 | 4.04 |
| 48 | 35.7 | 4.43 | 4.40 |

Determined based on a frictional ratio of 2.67. See Methods for details on the formula used for calculating the MW of a species with a given sedimentation coefficient and frictional ratio.

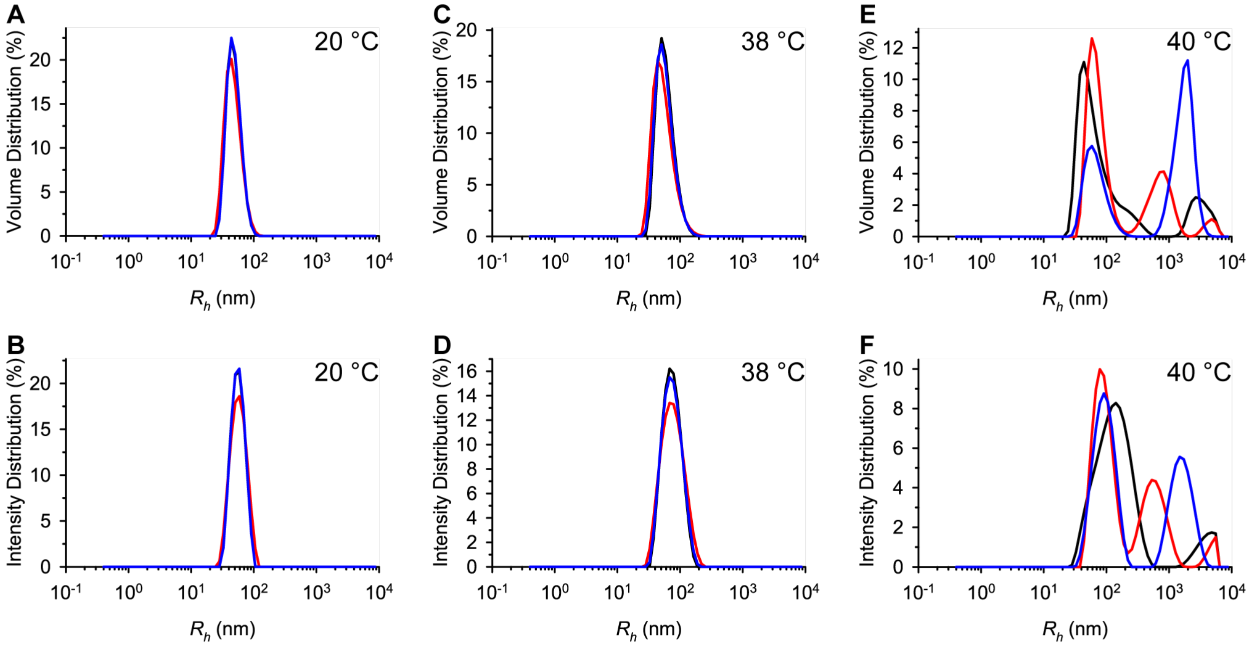


**Figure S7. DLS distributions of Brpt6.5 WT + $ZnCl_2$ at increasing temperature.** A sample of Brpt6.5 in the presence of 3.50 mM $ZnCl_2$ was measured by DLS. The data shown here are Volume and Intensity Distributions of the data shown in Figure 7B in the main text. Distributions are shown for data collected at 20 °C (A, B), 38 °C (C, D), and 40 °C (E, F). Resulting $R_h$ values are shown in Table S5.

**Table S5. DLS measurements of particle size.**

| Temperature | $R_h$ (nm) | Polydispersity Index |
|---|---|---|
| | 27.26 ± 0.16 | 0.033 |
| 30 °C | 27.34 ± 0.18 | 0.041 |
| 36 °C | 30.09 ± 0.05 | 0.047 |
| 38 °C | 34.93 ± 0.07 | 0.117 |
| 40 °C | 64.74 ± 3.07 | 0.499 |

Determined at 0.5 mg/ml Brpt6.5 WT in 50 mM MOPS pH 7.2, 50 mM NaCl, 3.50 mM $ZnCl_2$. Triplicate measurements were taken at each temperature to obtain an average $R_h$. Values in parenthesis indicate 1 standard deviation.
